# Unveiling hidden eukaryotes: diversity of Endomyxa (Rhizaria) in coastal marine habitats

**DOI:** 10.1101/2022.07.18.500052

**Authors:** Cédric Berney, Stefan Ciaghi, Sarah Romac, Frédéric Mahé, Colomban de Vargas, Olivier Jaillon, Martin Kirchmair, David Bass, Sigrid Neuhauser

**Author notes:** Correspondence: Sigrid Neuhauser, University of Innsbruck, Institute of Microbiology, Technikerstraße 25, 6020 Innsbruck, Austria, E-Mail, David Bass, Centre for Environment, Fisheries and Aquaculture Science (Cefas), Barrack Road, The Nothe, Weymouth, DT4 8UB, UK. Contributed equally to this work (Cédric Berney, Stefan Ciaghi).

## Abstract

Nucleic acid based studies of marine biodiversity often focus on Kingdom-level diversity. Such approaches often largely miss diversity of less studied groups, likely to harbour many unknown lineages which are likely playing significant ecological roles. Among these elusive groups are Endomyxa (Rhizaria), a ubiquitous, but understudied lineage comprising parasites (e.g. Ascetosporea, Phytomyxea), free-living amoebae (e.g. Vampyrellida, *Gromia, Filoreta*), flagellates (e.g. *Tremula, Aquavolon*), and unknown environmental lineages. Using Endomyxa-biased primers targeting the hypervariable region V4 of the 18S rRNA gene, we explored the diversity of Endomyxa in marine samples from European coastal sites and compared it to that found in pan-eukaryote V4 libraries of the same samples. In total 458 endomyxan OTUs were identified, of which 38% were only detected by the specific primers. Most are distinct from published sequences, and include novel diversity within known clades, and putative novel lineages. The data revealed variations in endomyxan assemblages related to habitat (benthic vs. pelagic, sampling site), mode of nutrition (parasitic vs. free-living) and nucleic acid type (DNA vs. RNA). Overall, the vast majority of endomyxan diversity occurs in sediments (including Vampyrellida, Reticulosida, and the environmental “Novel Clade 12”) where they form diverse and active communities including many uncharacterised lineages.

## Introduction

More than two thirds of the earth’s surface is covered by oceans, making them the largest biome on our planet. Therefore, analysing and understanding the diversity of marine microbes, including protists (Caron et al., 2009), and their roles in the ecosystem is a major interest of current research (Not et al., 2007; DeLong, 2009; Berney et al., 2013; Forster et al., 2016; Piredda et al., 2017; Tragin et al., 2018). Large projects like the European *BioMarKs* project (http://www.biomarks.eu/; Logares et al., 2014; Massana et al., 2015) and the *Tara* Oceans project (http://www.embl.de/tara-oceans/start/; Brum et al., 2015; de Vargas et al., 2015; Lima-Mendez et al., 2015; Sunagawa et al., 2015; Villar et al., 2015) aimed to investigate and describe marine microbial and viral communities using high throughput sequencing (HTS). Generally, the gene coding for the small subunit rRNA (18S rDNA) or its corresponding RNA is used for biodiversity and phylogenetic analyses of protists (Caron et al., 2012). The hypervariable regions V4 (Logares et al., 2012) and V9 (de Vargas et al., 2015) or both (Stoeck et al., 2010; Dunthorn et al., 2014; Logares et al., 2014; Massana et al., 2015; Tragin et al., 2018) of the 18S rDNA are the most widely used markers in environmental DNA (eDNA) studies (Pawlowski et al., 2012).

For several reasons, however, such metabarcoding approaches necessarily fail to uncover the full extent of eukaryotic diversity in any studied environment. The percentage of this diversity that can be retrieved is directly dependent on (1) the universality of the PCR primers used, and (2) size variation in the 18S rDNA fragment targeted by these primers - caused mainly by specific expansions in the most variable regions of the gene (e.g. Wuyts et al., 2001), or the presence of introns in the conserved regions of the gene (e.g. Bachar et al., 2013). As a result, eukaryotic lineages with more divergent and/or longer 18S rDNA sequences are likely to be systematically biased against during the PCR step of metabarcoding surveys (for examples of such lineage-specific size variation in the V4 region, see figure 5 in Wuyts et al. (2000)). Using lineage-specific primer sets is the best approach to explore the diversity within these lineages (e.g. Bass et al., 2012; Hartikainen et al., 2014a; Hartikainen et al., 2014b; Bass et al., 2015), although the specificity of the primers means that potential novel diversity around divergent lineages may still be missed.

Among the groups that belong to this “elusive” diversity of eukaryotes are the Endomyxa, a diverse group of protists belonging to Rhizaria (Bass et al., 2009) which often display divergent 18S rDNA sequences. Cavalier-Smith (2002) introduced Endomyxa to group together Ascetosporea and Phytomyxea, both rhizarian endoparasites of a wide range of organisms. Phytomyxea infect marine autotrophs, such as seagrasses (Sullivan et al., 2017), brown algae and diatoms (Schwelm et al., 2017), as well as many terrestrial angiosperms (Neuhauser et al., 2014). Ascetosporea are parasites of marine invertebrates (Hartikainen et al., 2014a; Hartikainen et al., 2014b; Ward et al., 2016; Ward et al., 2018). Gromiida, which are free-living, deep sea testate amoebae, also belong to Endomyxa (Cavalier-Smith and Chao, 2003a). Reticulosida, including the marine, network-forming amoebae of the genus *Filoreta*, were a later addition to that group (Bass et al., 2009). Also part of Endomyxa are the Vampyrellida (Hess et al., 2012; Berney et al., 2013), mainly known for their peculiar way of eating by drilling a hole in the cell walls of their host/prey (typically algae and fungi), although some species eat in a more typical “amoeba-like” way by engulfing whole prey cells. Although originally thought to be largely terrestrial and aquatic, vampyrellids have been shown to be very diverse in marine habitats (Berney et al., 2013). In addition to this known organismal biodiversity, several distinct, 18S rDNA-based eDNA lineages have been identified in Endomyxa (e.g. Bass and Cavalier-Smith, 2004; Bass et al., 2009).

In spite of their high ecological and evolutionary interest, Endomyxa remain understudied. Because of generally higher 18S rDNA divergence in endomyxan and retarian lineages compared to most of the closely related filosan lineages, most eukaryote-wide environmental surveys lead to a vision of rhizarian diversity that is overwhelmingly dominated by Filosa (=Cercozoa). In this study, we use Endomyxa as a case study to explore the potential of a mixed primer approach (combining group-specific and more general primers) to target the diversity within and around lineages that are typically missed in eukaryote-wide environmental surveys. We present the most comprehensive study so far on the diversity of Endomyxa in coastal marine habitats, focusing on three near-shore sites across Europe’s coastline. Samples from Naples, Oslo and Varna, collected as part of the *BioMarKs* project, were screened with Endomyxa-biased sets of primers spanning the V4 region of the 18S rDNA. Our results allow us to (i) assess the diversity of andomyxa (ii) identify differences in geographically (and therefore ecologically) separated endomyxan communities, (iii) identify differences between pelagic and benthic communities, and (iv) compare endomyxan biodiversity in DNA-versus RNA-derived samples.

## Results

### Endomyxa-biased versus Eukaryote-wide libraries

Although (the discontinued) 454 sequencing generated significantly lower read numbers than Illumina sequencing, the Endomyxa-biased 454-libraries resulted in a strikingly higher proportion of endomyxan reads (42,976 out of 210,126 [20.5%] versus 56,204 out of 9,656,192 [0.6%] Table 1, Figure S1). In addition, in the Endomyxa-biased datasets endomyxan reads were distributed more evenly across the known endomyxan lineages in comparison to endomyxan reads found in the eukaryote-wide libraries (Figure 1). In particular, the vast majority of endomyxan reads in the eukaryote-wide libraries belong to a single lineage, the vampyrellids (44,734 out of 56,204 [79.6%] versus 926 out of 42,976 [2.2%] in the Endomyxa-biased libraries). One consequence of the more even phylogenetic distribution of the Endomyxa-biased libraries was the exclusive detection of gromiids and ascetosporeans. Generally, the proportion of OTUs was also higher in the Endomyxa-biased dataset for all endomyxan lineages, except vampyrellids and phytomyxids (for which proportion of OTUs were roughly equivalent in both approaches). It is also noteworthy that the overlap of OTU richness between the two approaches is low.

**Table 1:**
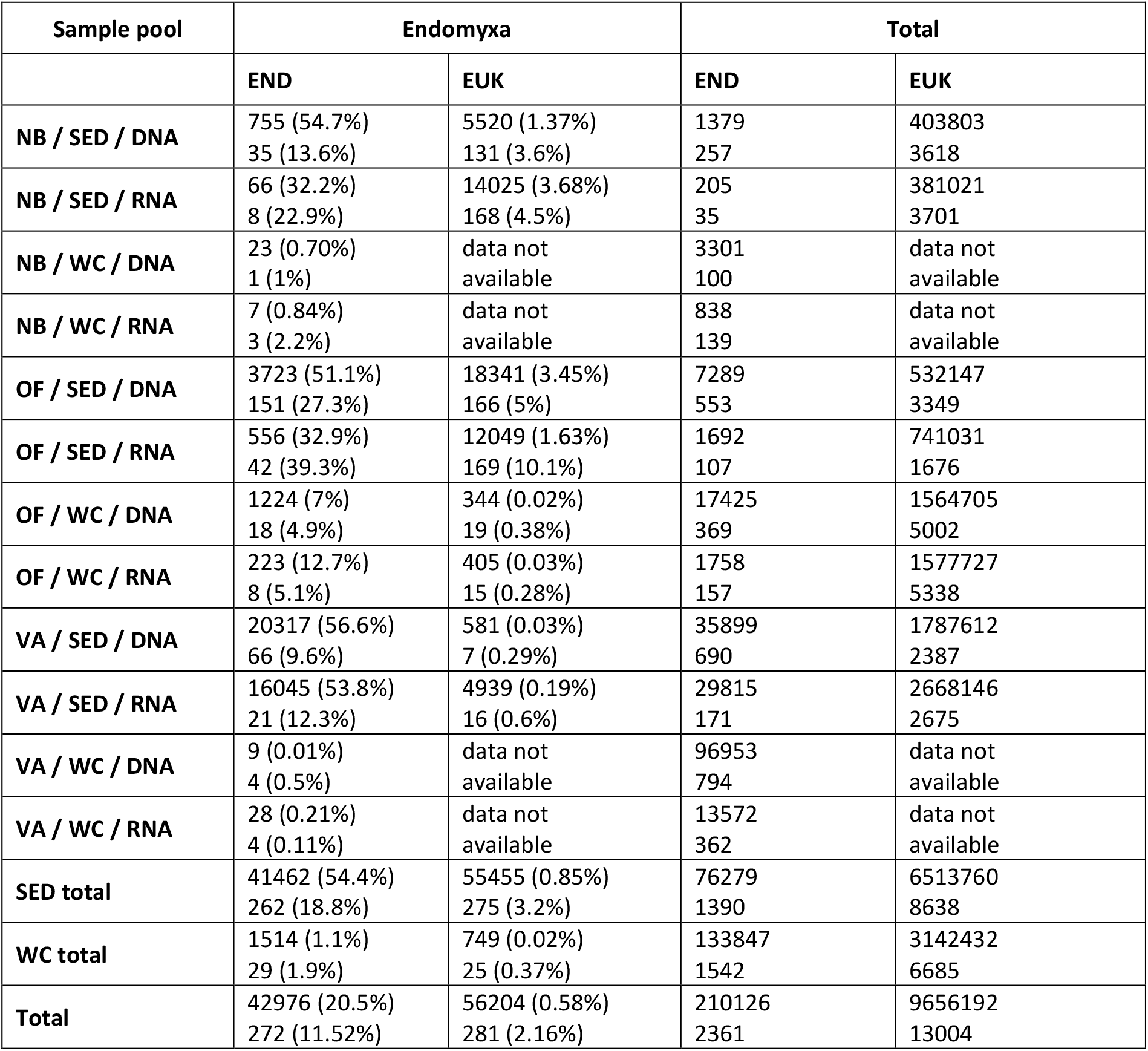
Endomyxa reads. Number and percentage of reads (first line) and OTUs (second line) identified in Endomyxa-biased libraries (‘END’, 454 sequencing) and eukaryote-wide libraries (‘EUK’, Illumina sequencing) from corresponding samples. ‘NB’: Naples Bay; ‘OF’: Oslofjord; ‘VA’: Varna; ‘SED’: sediment samples; ‘WC’: water column samples. Data not available means that no illumina data were generated from these samples.

**Figure 1:**
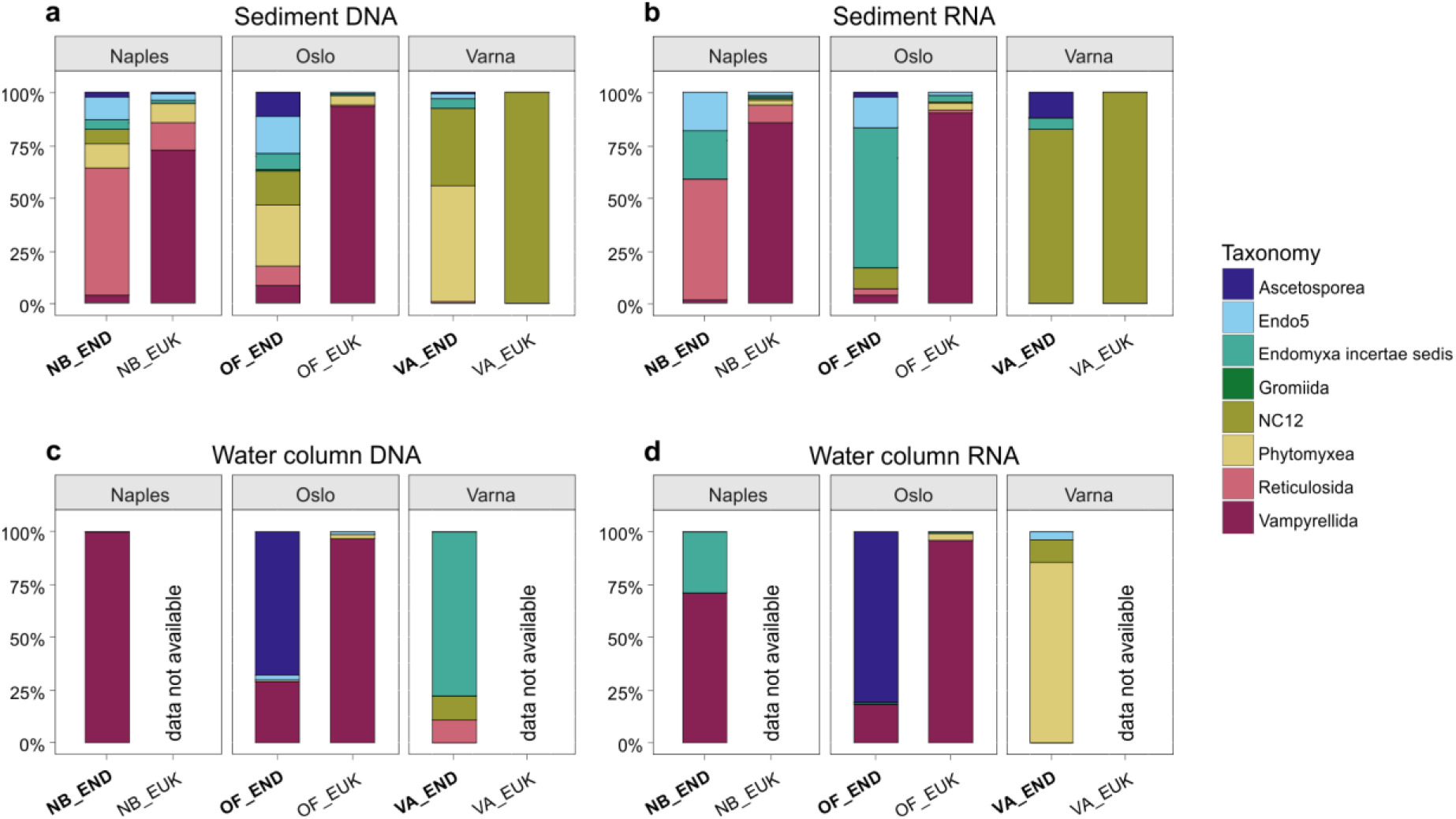
Relative abundance and diversity of Endomyxa reads in sediment DNA (a) and RNA (b) samples and water column DNA (c) and RNA (d) samples. Our 454-sequenced, Endomyxa biased libraries (‘END’; bold letters) were more diverse than the eukaryote-wide Illumina libraries (‘EUK’). The diversity in the water column was lower than in the sediment samples. Illumina libraries were not available for all water column samples. ‘NB’: Naples Bay; ‘OF’: Oslofjord; ‘VA’: Varna.

### Endomyxa OTU richness and phylogenetic diversity

In total, from both primer strategies, 458 OTUs could be taxonomically assigned to endomyxan lineages (Table S1). All known marine lineages of Endomyxa with morphologically described members were found in the dataset (Figure 2), with the exception of the three most divergent groups within Ascetosporea (Haplosporida, Mikrocytida, and Paramyxida). 204 OTUs were assigned to Vampyrellida, 31 to Phytomyxea, 27 to *Filoreta* spp. (Reticulosida), 26 to Ascetosporea, and 3 to Gromiida. Nearly a quarter of the OTUs (104) were assigned to Novel Clade 12, and 24 OTUs were assigned to lineage Endo5, both of them previously detected but still morphologically uncharacterised environmental lineages. Finally, 39 OTUs could be identified as endomyxans but could not be assigned to previously known lineages and were therefore labelled as “Endomyxa *incertae sedis*”. The phylogenetic tree shown in Figure 2 (restricted to Endomyxa) allows a better assessment of the phylogenetic diversity behind the OTUs labelled as “*incertae sedis*”. The tree topology is congruent with previous studies (e.g. Bass et al., 2018) and shows endomyxans separating into three major groups. The first one contains Vampyrellida and Phytomyxea; the second one contains Ascetosporea, Gromiida, Reticulosida, and environmental lineages Endo4 and Endo5; the third one contains Aquavolonida and Tremulida (two lineages found so far exclusively in non-marine habitats, and therefore absent in our libraries) as well as environmental Novel Clade 12 (found both in marine and terrestrial habitats). Three OTUs *incertae sedis* branch at the base of Vampyrellida (see also Figure S2), while five OTUs branch at the base of Ascetosporea + Gromiida and another five branch near lineages Endo4 and Endo5 (see also Figure S3). All other OTUs *incertae sedis* branch within the Aquavolonida + Tremulida + Novel Clade 12 group (see also Figure S4) and further support the idea that unlike the well-defined Aquavolonida and Tremulida, “Novel Clade 12” actually represents a poorly resolved radiation of many marine and/or terrestrial lineages.

**Figure 2:**
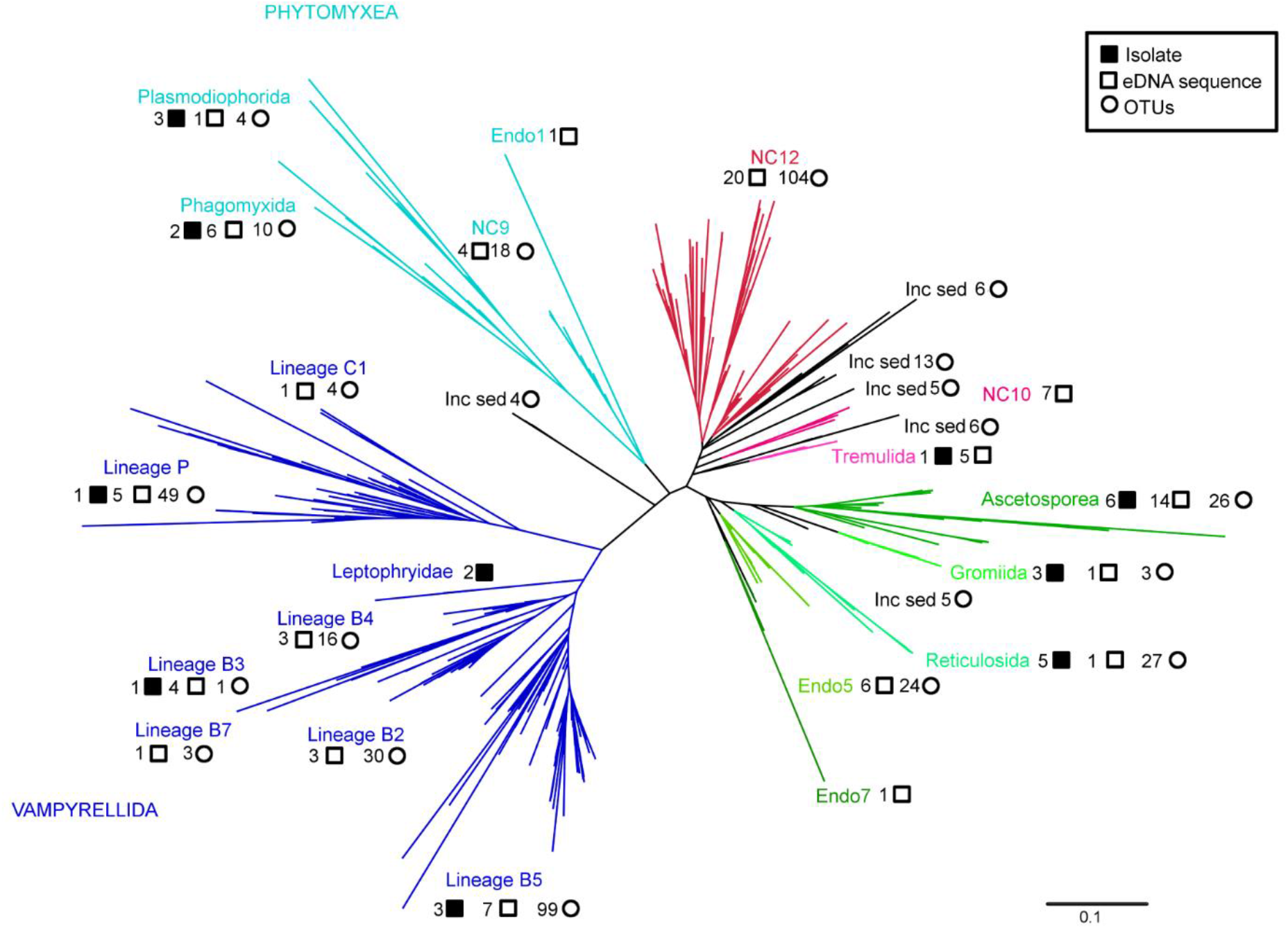
Phylogenetic placement tree of marine Endomyxa OTUs. Clades which had been described previously are given in colour, novel lineages (“inc sed”) are given in blue. Black, filled squares highlight sequences that originate from isolates, open squares indicate eDNA sequences previously used in phylogenetic studies, circles are OTUs found in this study. More detailed phylogenetic trees can be found in the Supplement (Figures S2-S5).

### Comparisons between sites, habitat pools, and templates

The vast majority of endomyxan OTUs (410 out of 458) were found only in sediment samples, with the exception of 10 OTUs which were found in the water column only, and 38 were found in both sample types (Table S2). The relative abundance of endomyxan reads was also higher in sediment samples than in water column samples (Table S2). Vampyrellids were the most commonly detected group in water column samples (in Naples the only one), but ascetosporean OTUs were detected in fairly high relative abundance in the Oslo water column samples. In the sediment, vampyrellids clearly dominate the Naples and Oslo samples both in terms of OTU richness and relative read abundance in the eukaryote-wide libraries (Figure S5). However, vampyrellid relative abundance drops significantly in the Endomyxa-biased libraries, where many groups were detected at comparable relative abundance and OTU richness both in Naples and Oslo (Ascetosporea, Reticulosida, Phytomyxea, and environmental Novel Clade 12 and lineage Endo5, Table S2). By contrast, the Varna sediment samples are dominated by Novel Clade 12 (the only lineage detected there in the eukaryote-wide libraries), and this was observed in both DNA- and RNA-derived samples (Figures 3 and S6). Phytomyxea (specifically an environmental lineage within them referred to as Novel Clade 9) was also detected at a fairly high proportion, but only in the DNA-derived samples (Figures S5, S7, S8). Both incidence-based and abundance-based ordinations of the endomyxan communities found in the six sample pools (Figure 4) confirm that the Varna samples (Black Sea) have significantly distinct communities compared to the Naples and Oslo samples; among the latter samples are primarily segregated based on habitat type (sediments versus water column). In the Endomyxa-biased libraries, on average across all sites and in both habitat types, fewer endomyxan reads were detected in the RNA-derived samples (Figure 3, S5, S9, S10). However, these systematically contained fewer usable reads in total (Table S2).

**Figure 3:**
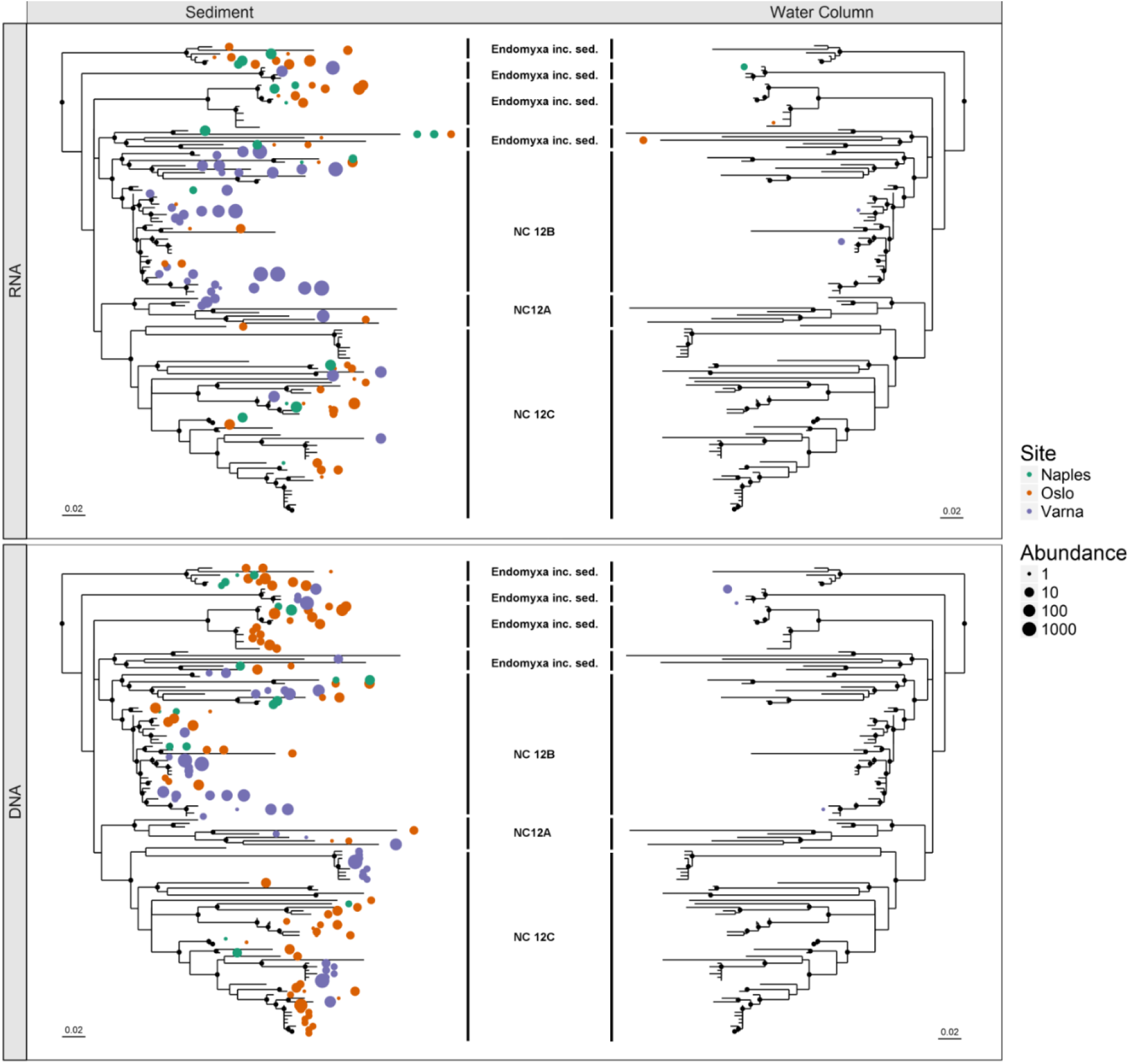
Abundance (read number) and frequency (libraries where they were detected) of OTUs belonging to Novel Clade 12 (NC12). NC12 was predominantly found in sediment samples and NC12 OTUs are highly abundant in the Varna RNA samples. Each sample that contained a respective OTU is shown as an individual dot, while the size indicates the number of reads that were recovered from each library. Black dots at the branching points indicate χ^2^ values above 0.95 from the PhyML analyses. Trees were generated using the R-package ‘phyloseq’.

**Figure 4:**
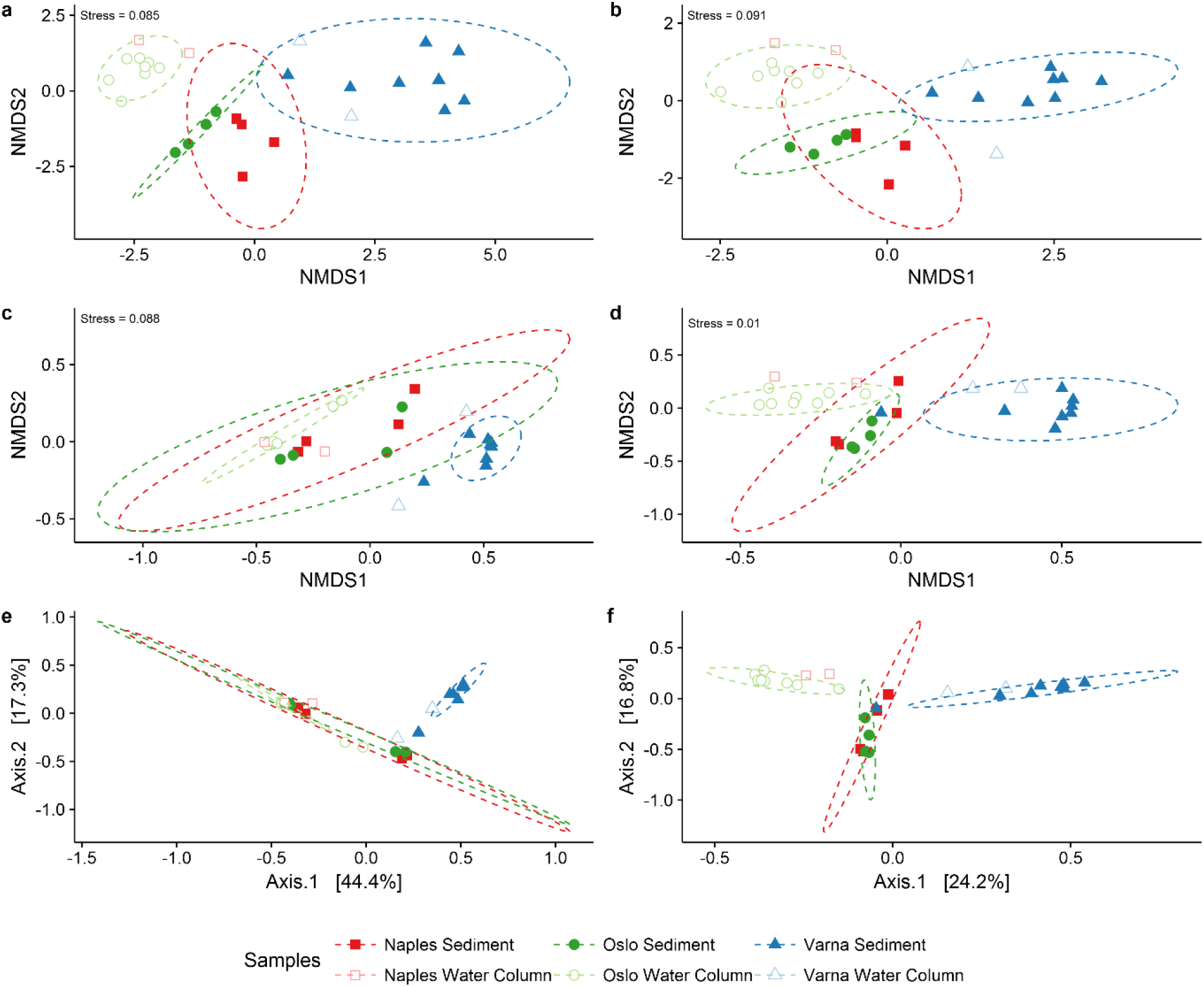
NMDS and PCoA sample patterns based on abundance (a,c,e) and incidence (b,d,f) data. The Varna samples were clearly separated from the other samples in all plots. A difference between water column and sediment samples was observed, particularly looking at Samples from Naples and Oslo. a: NMDS ordination using Bray-Curtis dissimilarity. b: NMDS ordination using Jaccard dissimilarity. c: NMDS ordination using weighted UniFrac distance. d: NMDS ordination using unweighted UniFrac distance. e: PCoA ordination using weighted UniFrac distance. f: PCoA ordination using unweighted UniFrac distance. Confidence ellipses (95%) are given where enough samples for calculation were present.

### Endomyxa-biased PCR primers and libraries

We designed an original nested PCR approach based on a combination of several specific and universal primers (Table S3) to cover as much of the total diversity of Endomyxa as possible (but necessarily excluding the most extremely divergent ascetosporeans: mikrocytids and paramyxids). In the first PCR, we used a universal forward primer upstream of variable region V4 with a pool of four reverse primers at the 3’ end of the 18S rDNA designed to cover the known diversity within most endomyxan lineages, and specifically excluding Filosa, which display two rare substitutions in the descending part of the helix corresponding to variable region V9 (Cavalier-Smith and Chao, 2003a). In the nested PCR, two forward primers near the beginning of variable region V4 were pooled; one covering most eukaryotes and the other ending on a substitution present in all endomyxans but absent in most other eukaryotes (Bass et al., 2018). These were used with a pool of 10 reverse primers situated between variable regions V5 and V7 and spanning a rare, one nucleotide deletion specific of Endomyxa and Filosa (Cavalier-Smith and Chao, 2003b). Several versions of the primer were pooled to cover most known sequence variants in endomyxans, yet without using a single, too highly degenerate primer. This primer pool was biased towards a version containing no ambiguity and representing the consensus sequence found in Endomyxa (s1256R-C0) in order to favour detection of potentially rare, but non-divergent endomyxan lineages.

In total, the 454-sequenced Endomyxa-biased libraries from the BioMarks Project resulted in 210,126 reads that successfully passed our quality checks and were long enough to cover the full V4 fragment and be comparable with the reads obtained from the available corresponding samples in the Illumina sequenced, eukaryote-wide libraries (which by contrast included a total of 6,513,760 reads). The 454-sequenced reads that passed our quality checks were then trimmed to the length corresponding to the fragment sequenced in the eukaryote-wide libraries, and pooled with the Illumina-sequenced reads for OTU clustering (see Material and Methods).

## Discussion

### Targeting Endomyxa necessitates a specific primer strategy

Endomyxa are a genetically divergent group; this makes them extremely difficult to study as a whole using data from eukaryote-wide eDNA studies, because of length polymorphism and variations in their 18S rDNA sequences (Bass et al., 2009; Sierra et al., 2016). Existing eDNA surveys and HTS datasets based on universal V4 primers typically harbour a very low diversity of endomyxans, with only two notable exceptions. In freshwater habitats, the recently characterised aquavolonids (“Novel Clade 10”) are frequently detected in the water column (Bass et al., 2018 and references therein). In soils and in marine sediments, the same is true for vampyrellids (Berney et al., 2013; Massana et al., 2015). Both of these lineages have “normal” 18S rDNA sequences matching typical eukaryote-wide primers and lacking specific extensions, and they appear abundant enough (in the specific habitat types mentioned above) to be recovered at a significant level compared to other eukaryotic groups. In the case of vampyrellids, this is also supported by their frequent observation in mixed cultures from soils and coastal sediment samples (Berney et al., 2013). By contrast, other endomyxan lineages are only rarely detected. For some of them, it could be that they are truly rare or confined to specific, understudied habitats (see below); in several cases however, it can be expected to be a result of their divergent 18S rDNA sequences. For these, using highly specific PCR primer approaches has proven efficient to uncover unknown diversity within individual lineages such as haplosporidians (Hartikainen et al., 2014a), mikrocytids (Hartikainen et al., 2014b), paramyxids (Ward et al., 2016), or phytomyxeans (Neuhauser et al., 2014). However, these approaches are so specific that it is unlikely they could reveal undetected endomyxan diversity outside of the targeted lineage.

In this study, existing data from a mixed primer approach was analysed to allow a more exhaustive detection of the full diversity of Endomyxa in marine habitats. The aim of this approach was to target rhizarian diversity broadly enough to uncover potential new endomyxan lineages, while at the same time excluding as much as possible of the diversity of non-endomyxan Rhizaria, especially Filosa (see Results for details of the specificity levels of the primers we used).

### Endomyxa global diversity and prevalence

Our mixed PCR approach enabled what we believe to be the most exhaustive coverage to date of the total diversity of Endomyxa in marine coastal habitats. The relative abundance of Endomyxa reads compared to all other eukaryotes was much higher in the Endomyxa-biased libraries (ca. 20%) versus the eukaryote-wide libraries (less than 1%), indicating that the primers we used successfully favoured amplification of endomyxans. Importantly, we detected endomyxan OTUs that could not be assigned to known Endomyxa groups and potentially represent new lineages in that major rhizarian clade (Figure 2, Figure S1-S4). This would not have been possible with lineage-specific primers. Furthermore, the striking differences in OTU richness and relative abundance of the various endomyxan lineages in the Endomyxa-biased versus eukaryote-wide libraries confirm that beyond a certain point, higher sequencing depth alone is not enough to improve detection of the total diversity of certain eukaryotic groups (Figure S1). Even though our Endomyxa-biased libraries contain significantly lower amounts of usable reads because of the technical limitations of using a dataset generated with the outdated 454 platform the overall diversity they uncovered was higher, still yielding valuable and novel data on the diversity of this group. In particular, two groups (Ascetosporea and gromiids) that contain specific insertions in the V4 region which are therefore not detected with eukaryote-wide primers could be detected in the Endomyxa-biased libraries (Figure 1). Overall, the two most diverse endomyxan lineages in marine coastal habitats seem to be the vampyrellid amoebae (as shown by Berney et al., 2013), Figure 2, Figure S2) and Novel Clade 12 (which was never highlighted before, Figure 3, Figure S4). Other significantly diverse groups include Phytomyxea (parasites of algae, Figure S7), Ascetosporea (parasites of invertebrates), Reticulosida (large, naked reticulose amoebae), and environmental lineage Endo5 (of unknown phenotype, Figure S3).

However, it is important to note that the relative abundances of the various endomyxan lineages uncovered in the Endomyxa-biased libraries are not necessarily more comprehensive than those observed with eukaryote-wide primers. First, even though our mixed primers approach covers most of the known diversity of Endomyxa, at least three divergent ascetosporean lineages (Haplosporida, Mikrocytida, and Paramyxida) cannot be amplified with our primers (Table 1, Figure S4). Lineage-specific primers remain necessary to access the diversity of these divergent taxa, making it impossible to directly compare their prevalence to that of other endomyxan lineages. Second, we specifically designed our primer approach to favour detection of potentially rare Endomyxa with non-divergent 18S rDNA sequences. We believe this explains the much lower relative abundance of vampyrellids in Endomyxa-biased libraries compared to eukaryote-wide libraries (Table 1). At least in the case of lineages with normal 18S rDNA sequences that do amplify with eukaryote-wide primers (these would notably include Reticulosida and environmental Novel Clade 12 and Endo5) their relative abundances to other taxa suggested by eukaryote-wide libraries are probably more realistic. Therefore, we can assume that among endomyxans successfully covered by eukaryote-wide primers, vampyrellids largely dominate in the Naples and Oslo sediment samples, while Novel Clade 12 is the most diverse and abundant lineage in the Varna samples.

### Distribution of marine Endomyxa

An overwhelming majority of the endomyxan diversity we uncovered (98% of OTUs) was found in the benthos, as opposed to only 10% of OTUs found in the water column (Figure 1, 3, Figure S5, S8). Sediments are a logical habitat for large reticulose/ramose amoebae like Vampyrellida and Reticulosida (*Filoreta* spp.). For such amoeboid predators, this morphotype is expected to allow optimal exploration of a granulated environment in search of suitable prey (as discussed in Berney et al., 2015). It can be hypothesised that parasitic lineages like Ascetosporea and Phytomyxea will more likely be found in the sediments than in the water column, either because their hosts have a benthic lifestyle, or because sediments act as a reservoir of dormant spores (e.g. Chambouvet et al., 2014). Whenever these do occur in the water column (as observed in the Oslo samples for Ascetosporea, Figure S10), we can expect that they correspond to active, infective stages. However, they may also represent actual infections of planktonic hosts like crustaceans (e.g. paradinids) or planktonic algae and stramenopiles (various phytomyxids).

As for the distinctiveness of the Varna sediment sample pools compared to the Naples and Oslo samples, it can be explained by the fact that they contained samples from the anoxic/low oxygen environments. In these, the free-living amoebae dominating the Oslo and Naples samples were hardly detected, while Novel Clade 12 and Endomyxa *incertae sedis* representing new lineages closley related to Novel Clade 12 showed a high diversity and relative abundance (Figure 3, S6). This result points at oxygen-depleted environments as key habitats to find previously undetected eukaryotic diversity.

### Environmental Novel Clade 12 and Endo5 - what are they?

A major finding of this study is the realisation that the few previously known environmental clone sequences that phylogenetically grouped together into what was called Novel Clade 12 (NC12) only represented the tip of the iceberg of a much more substantial radiation of 18S rDNA lineages (Figure 3, Figure S4). NC12 was first detected in Bass et al. (2009), where it was shown to comprise environmental sequences from mostly anaerobic/anoxic habitats, both terrestrial and marine (including deep sea). NC12 lineages are phylogenetically related to two groups (labelled Novel Clades 10 and 11 in Bass et al. (2009)) that have since been morphologically characterized as Aquavolonida (Bass et al., 2018) and Tremulida (Howe et al., 2011), respectively. In addition to widely expanding the diversity around previously known NC12 environmental clones, our new data revealed OTUs (labelled Endomyxa *incertae sedis*) that are phylogenetically as distant from tremulids and aquavolonids as they are from “true” NC12 clones (Figure S4). Existing studies (e.g. Bass et al., 2018) suggest that together, they occupy a key phylogenetic position within Rhizaria, branching deeply in the supergroup, separate from both Filosa (= core Cercozoa) and the rest of Endomyxa (all the groups displayed in green and blue tones in Figure 2). In this context, morphological characterization of members of this “NC12 radiation” is important to infer the ancestral phenotype in Rhizaria. Unlike most other Rhizaria, tremulids and aquavolonids are eukaryovorous flagellates without amoeboid tendencies. The most likely hypothesis is that the various lineages comprising the “NC12 radiation” also contain “simple” flagellates resembling the genera *Tremula* and *Aquavolon*; i.e. that they may be eukaryovorous flagellates. The OTUs belonging to the “NC12 radiation” detected in this study seem to be particularly diverse in oxygen-depleted sediments, and present both in marine and terrestrial habitats. Tremulids and aquavolonids are restricted to terrestrial habitats, and aquavolonids in particular show an adaptation towards a freshwater planktonic lifestyle (Bass et al., 2018). However, the Aquavolonida + Tremulida + NC12 clade as a whole can be hypothesized to have originated in oxic, benthic, marine habitats. Competition from other, more successful phenotypes (including the highly diverse core Cercozoa) may have pushed Aquavolonida to adapt to a planktonic lifestyle and NC12 to diversify in anoxic sediments.

Diversity within another environmental lineage Endo5 is here also revealed as much higher than previously thought (Figure 2, Figure S3). Endo5 was also first detected by Bass et al. (2009) based on a single partial 18S rDNA clone from oxic marine sediments (sm5, GenBank accession number EU567281). Since then, the lineage was only detected again three times based on GenBank searches for endomyxan clone sequences (clone RA070411T.004 [FJ431660] from Marie et al. (2010), clone BS19_B6 [FN598385] from Sauvadet et al. (2010), and clone 34c_10165 [KT815328] from Marquardt et al. (2016)). All of these clones only span part of the 18S rDNA, which is why we generated two full-length 18S rDNA clone sequences from environmental DNA extractions corresponding to two distinct Endo5 phylotypes (see Material & Methods) to improve robustness of the phylogenetic placement of that lineage. Endo5 belongs to the endomyxan clade that comprises Reticulosida, Gromiida, Ascetosporea, and another, even less frequently detected environmental lineage (Endo4 in Bass et al., 2009). Even though Endo5 seems significantly less diverse and abundant than the “NC12 radiation”, it nonetheless probably represents a relatively deep radiation within Endomyxa (our two new full-length sequences share only ca. 90% overall identity). The likely phenotype of Endo5 members is more difficult to infer than that of NC12 members because their closest characterized relatives include groups with very contrasting morphologies and lifestyles (the naked, reticulose *Filoreta* spp., the testate gromiids, and the ascetosporean parasites). Given their elusive nature, it is possible that Endo5 members are small (amoebo)flagellates.

### Conclusions and perspectives

Endomyxa as a group are difficult to study, as its members have contrasting modes of nutrition (predatory, osmotrophic, parasitic), are highly divergent in the 18S rDNA, and are in the best case difficult to isolate and maintain in culture. Therefore, eDNA or eRNA based studies are currently the only feasible way of efficiently approaching their ecology and biodiversity. Here we showed that using targeted primer sets greatly improved the taxonomic sampling within Endomyxa and revealed a significant diversity of previously undetected OTUs. This approach allowed us to detect novel lineages even within the parasitic groups that are most typically missed in eukaryote-wide studies. Apart from Vampyrellida in sediments (and possibly Novel Clade 12 locally in anoxic habitats), it remains unlikely that marine Endomyxa are ever particularly abundant compared to other eukaryotes. However the levels of diversity uncovered across Endomyxa when studied with appropriate primers (especially in sediment samples) would not be maintained if they did not play important ecological roles and/or did not have significant interactions with other members of their communities. The same situation can be expected to be true in other “difficult” eukaryotic groups, most of which have not yet been given any proper attention. With the constant improvement of sequencing technologies, including those that do not require a sequence-specific PCR step, the technical limitations we faced in this study because of the size of our Endomyxa-biased amplicons will soon no longer be an issue. In the meantime, the amount of novel biodiversity we found even in our extremely limited (in terms of samples, sequencing depth, and habitats) pooled libraries is proof of concept that approaches built on a targeted, but flexible primer strategy are a very powerful tool for future studies of elusive eukaryotes such as Endomyxa. Of immediate interest would be a endomyxan-focused eD/RNA study of non-marine habitats, particularly different soil types and freshwater habitats.

## Experimental Procedures

### Data

The raw data used for this study were generated as part of the European *BioMarKs* project (http://www.biomarks.eu/). In this study a sub-set of the samples originating from Naples (Tyrrhenian Sea, Italy, sampling 2009, 2010), Oslo (Oslofjorden, Norway, sampling 2009, 2010), and Varna (Black Sea, Bulgaria, 2010) were analysed (Table S4). Samples were collected from the sediment and from two different depths in the water column: subsurface and deep chlorophyll maximum (DCM); additionally, in Varna the anoxic layer at a depth of 250 m was also sampled. Detailed information about sampling and nucleic acid extraction and manipulation can be found in Logares et al. (2014), Massana et al. (2015), and on the website of the *BioMarKs* project (http://www.biomarks.eu/).

To analyse Endomyxa diversity, a targeted approach with Endomyxa-biased PCR primers was designed (Table S3; see Results for more details) and the resulting libraries were sequenced using the 454-pyro-sequencing platform (Roche, Branford, CT, USA). Samples were pooled via protags into “water column” samples (combining subsurface and DCM samples) and “sediment” samples, which in Varna included the 250 m anoxic layer samples (Table S4). These Endomyxa-biased libraries were compared to data generated by the *BioMarKs* consortium from a subset of the same samples amplified with the universal eukaryotic primers from Logares et al. (2014) and sequenced using the Illumina GAIIx sequencing platform (Illumina, San Diego, CA, USA).

### Endomyxa-specific libraries construction and sequencing

PCR products were generated from DNA and cDNA samples from Naples, Oslo, and Varna (Table S4) using a nested PCR approach. The DNAs and cDNAs were diluted to a concentration of 2 µg µL^-1^ and 4 µg DNA from each sample were used in the first round PCRs. These were done in 30 µL in a mix containing 6 mM dNTPs, 1.5 U GoTaq polymerase, 7.5 mM MgCl_2,_ 1x PCR buffer and 12 mM of each primer. First round PCR conditions were 95 °C for 3 min followed by 36 cycles of 95 °C for 30 s, 66 °C for 30 s, and extension at 72 °C for 90 s, followed by 10 min at 72 °C. The samples were then stored at 6 °C until further processing. Nested PCRs were done in 20 µL with the same PCR mix scaled down to this volume and using as template 2 µL from the corresponding first round PCRs. Nested PCR conditions were the same as above, but with an annealing temperature of 67.5 °C and 39 cycles Endomyxa-biased primers (Table S3) were used in combination with multiplexing tags (MIDs) to allow for a pooling of samples. Three different MIDs were used, allowing four pools of samples (one without MID); they were added to the forward primers used in the nested PCRs. All nested PCRs were done in triplicates, then pooled and concentrated using Amicon Ultra 0.5 ml 50 kDa spin columns (Merck, Darmstadt, Germany) according to the manufacturer’s instructions and loaded onto a gel. Bands within the expected size range (700-900 bp) were cut and cleaned using the Quiagen QIAquick Gel Extraction Kit (Qiagen, Hilden, Germany) following the manufacturer’s instructions. Extra care was taken to include the areas above the bright band at 700/750 bp, because some Endomyxa lineages are known to have insertions in the V4 region.

The four pools of samples were prepared for pyrosequencing according to the manufacturer’s instructions. For each about 1 μg of amplicons were sent to Genoscope (Évry, France) where they were processed and pyrosequenced on a 454 GS FLX Titanium system (454 Life Sciences, USA). The sample pools from Naples and Oslo were sequenced on a quarter of a 454-plate each, while the sample pool from Varna was sequenced on half a plate (Table S5).

### Sequence Processing

The FNA and QUAL files obtained after 454-sequencing of our 24 Endomyxa-specific samples were merged into FASTQ format using the command fastaqual_to_fastq of seq_crumbs v0.1.9 (https://bioinf.comav.upv.es/seq_crumbs/). Quality value offset was verified with vsearch v1.11.1 (Rognes et al., 2016). The amplicons generated with the Endomyxa-biased primers are longer than the amplicons of the eukaryote-wide *BioMarKs* dataset with which we wanted to compare them because they use different reverse primers. As a result prior to OTU clustering all reads from our Endomyxa-biased libraries had to be trimmed down to the length of the eukaryote-wide dataset using the sequence of the universal reverse primer (5’-ACTTTCGTTCTTGATYRA-3’). This was done using cutadapt v1.10 (Martin, 2011), which also removed the primer sequences from the reads. Sequence length and expected error were computed for all trimmed sequences before converting them into FASTA format, and dereplicating them using vsearch. The raw sequences from 155 published *BioMarKs* samples (454 sequenced) were re-analysed following the same *in silico* protocol.

The resulting FASTA file was clustered into OTUs using Swarm v2.1.8 (Mahe et al., 2014, 2015) with the option fastidious. Representative sequences for the obtained OTUs were checked for chimera using the uchime *de novo* algorithm (Edgar et al., 2011) as implemented in vsearch, and received a taxonomic assignment using the July 2016 release of the reference database PR^2^ (Guillou et al., 2013) and the method stampa (https://github.com/frederic-mahe/stampa).

### Alignments and Trees

To make sure that all OTUs assigned to Endomyxa represent believable endomyxan diversity, we analysed them in the context of a manually curated, full length 18S rDNA alignment of 120 reference Endomyxa sequences, including 32 described species or observed isolates, and 88 environmental clones (for their accession numbers see Table S6). In the case of the environmental lineage Endo5 for which no full-length 18S rDNA sequence was yet available, two new reference environmental sequences were generated. For this, we relied on the results of lineage-specific surveys of endomyxan diversity in various environments that were conducted at the NHM in London as part of a research project on Endomyxa (to be published elsewhere). Two marine sediment DNA samples that were found to contain two distinct Endo5 phylotypes were used to generate full-length 18S rDNA sequence with a combination of internal Endo5-specific primers and universal primers at the beginning and end of the 18S rDNA (Table S6).

A representative sequence of each putative endomyxan OTU was extracted from the dataset and automatically aligned with all others using MAFFT (Katoh and Standley, 2013) implemented in Geneious R9.1.5 (http://www.geneious.com; Kearse et al., 2012) using the default settings and a 1PAM/k=2 scoring matrix. The aligned OTUs were then incorporated into the reference Endomyxa alignment, which was improved manually to take the more variable regions into account. To phylogenetically place the OTUs, lineage-specific insertions and the most highly variable regions were masked. The final alignment contained 584 sequences and 1 500 characters. Trees were then generated using PhyML v3.0 (Guindon et al., 2010) and χ^2^ support values and RAxML v7.2.8 (Stamatakis, 2014). Sub-trees of the PhyML tree were extracted as needed to conduct further analyses.

### Statistical analyses

Statistical analyses were performed in R v3.3.2 (R Core Team, 2016) using the ‘phyloseq’ package v1.18.1 (McMurdie and Holmes, 2013). The data were sub-sampled for Endomyxa OTUs only. Absolute and relative abundance analyses were performed to investigate the distribution of reads across the different samples. Distance based methods (Bray-Curtis dissimilarity, Jaccard dissimilarity, and UniFrac distance (Lozupone and Knight, 2005)) were used to measure β-diversity. Weighted UniFrac distance was calculated using total branch length normalisation. Non-metrical multidimensional scaling (NMDS; Kruskal, 1964) and principal coordinate analysis (Gower, 1966) were used to create ordination plots. The auto-transform option of the function metaMDS for calculating NMDS with the R package ‘vegan’ v2.4-1 (Oksanen et al., 2016), implemented in the ‘phyloseq’ package, was set for abundance based methods. This led to a square root transformation and Wisconsin double standardisation. Stress coefficients (Clarke, 1993) for NMDS ordinations were determined. Confidence ellipses (95%) were calculated based on the t-distribution. Bar plots for comparing the two datasets analysed using different primer strategies were created using the R package ‘ggplot2’ v2.2.1 (Wickham, 2009).

## Acknowledgements

We thank the Biodiversity of Marine euKaryotes consortium (*BioMarKs*; http://www.biomarks.eu), which was funded by the European Union ERA-Net program BiodivERsA (2008-6530). SN and SC were funded by the Austrian Science Fund: grant J3175-B20 (SN) and grant Y0801-B16 (SN, SC). This work was also supported by NERC grants NE/H009426/1 (DB, CB) and NE/H000887/1 (DB) and the ‘Investissements d’Avenir’ programme OCEANOMICS (ANR-11-BTBR-0008) awarded by the French Government via Agence Nationale de la Recherche (CB, SR, CdV). CB and CdV are also grateful to the Gordon and Betty Moore Foundation and the International Society of Protistologists for current funding (grant GBMF5257 / UniEuk project, www.unieuk.org).

