## Supplementary material for "Unveiling hidden eukaryotes: diversity of Endomyxa (Rhizaria) in coastal marine habitats": Figure S1 - Figure S10

***Unveiling hidden eukaryotes: the elusive Endomyxa (Rhizaria) are rich in function and diversity in coastal marine habitats***

***Supplementary Information - Figures***

Stefan Ciaghi[1,$], Cédric Berney[2,$], Sarah Romac[2], Frédéric Mahé[3], Colomban de Vargas[2], Olivier Jaillon[4], Martin Kirchmair[1], David Bass[5,6,*], Sigrid Neuhauser[1,*]

[1] Institute of Microbiology, University of Innsbruck, Technikerstr. 25, 6020 Innsbruck, Austria

[2] Sorbonne Université & CNRS, UMR 7144 (AD2M), Station Biologique de Roscoff, Place Georges Teissier, 29680 Roscoff, France

[3] CIRAD, UMR LSTM, Campus international de Baillarguet, 34398 Montpellier Cedex 5, France

[4] GENOSCOPE, Commissariat à l'Energie Atomique et aux Energies Alternatives (CEA), Institut de Génomique, 2 rue Gaston Crémieux, 91000 Evry, France

[5] Division of Genomics and Microbial Diversity, Dept of Life Sciences, Natural History Museum London, Cromwell Road, SW7 5BD, UK

[6] Centre for Environment, Fisheries and Aquaculture Science (Cefas), Barrack Road, The Nothe, Weymouth, DT4 8UB, UK

**Supplementary Figures S1- S10**


| | |
|---|---|
| **Figure S1:** Relative abundance of OTUs with Eukaryote and Endomyxa Primers | **2** |
| **Figure S2:** Tree - Biodiversity Vampyrellida | **Separate file** |
| **Figure S3:** Tree - Biodiversity Ascetosporea, Reticulosida, Gromiida, Endo5 | **Separate file** |
| **Figure S4:** Tree - Biodiversity Aquavolon, NC12, Tremula | **Separate file** |
| **Figure S5**: Abundance and frequency of Vampyrellida, Phytomyxea | **TBA** |
| **Figure S6:** Incidence tree NC12 | **TBA** |
| **Figure S7:** Tree - Biodiversity Phytomyxea | **seperate file** |
| **Figure S8**: Incidence tree Vampyrellida, Phytomyxea | **TBA** |
| **Figure S9:** Abundance tree Ascetosporea, Reticulosida, Gromiida, Endo5 | **TBA** |
| **Figure S10:** Incidence tree Ascetosporea, Reticulosida, Gromiida, Endo5 | **TBA** |
| **References** | **TBA** |

### Supplementary Figures

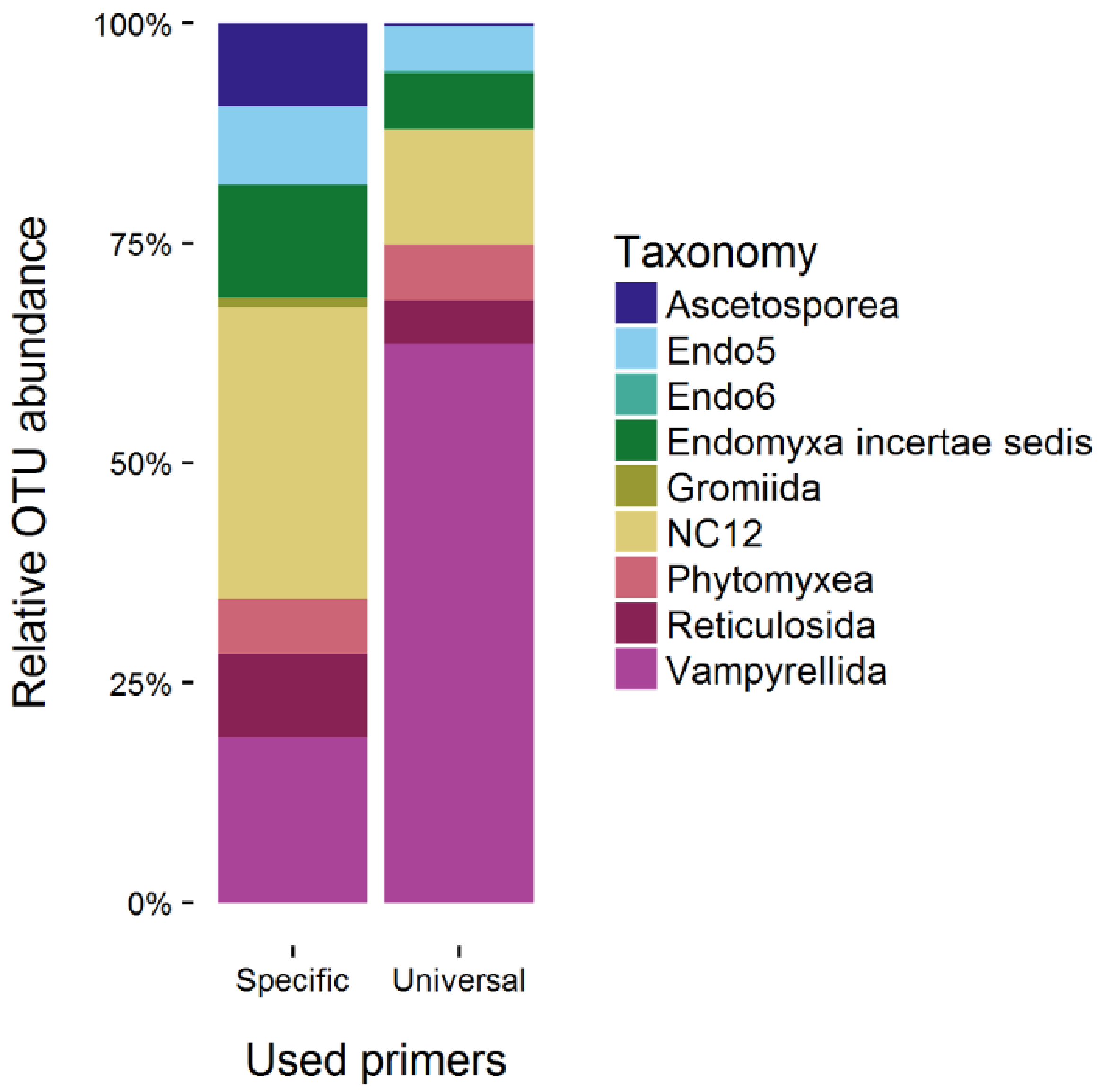


**Figure S1: Relative abundance of Endomyxa OTUs** in Endomyxa-specific (n = 2 263, 454) and pan-eukaryote (n = 96 917 Illumina) libraries. Despite the lower read number the biodiversity recovered from the 454 dataset with Endomyxa-specific primers was higher and more evenly spread across the taxonomic groups than when using a more generic approach with eukaryote primers.

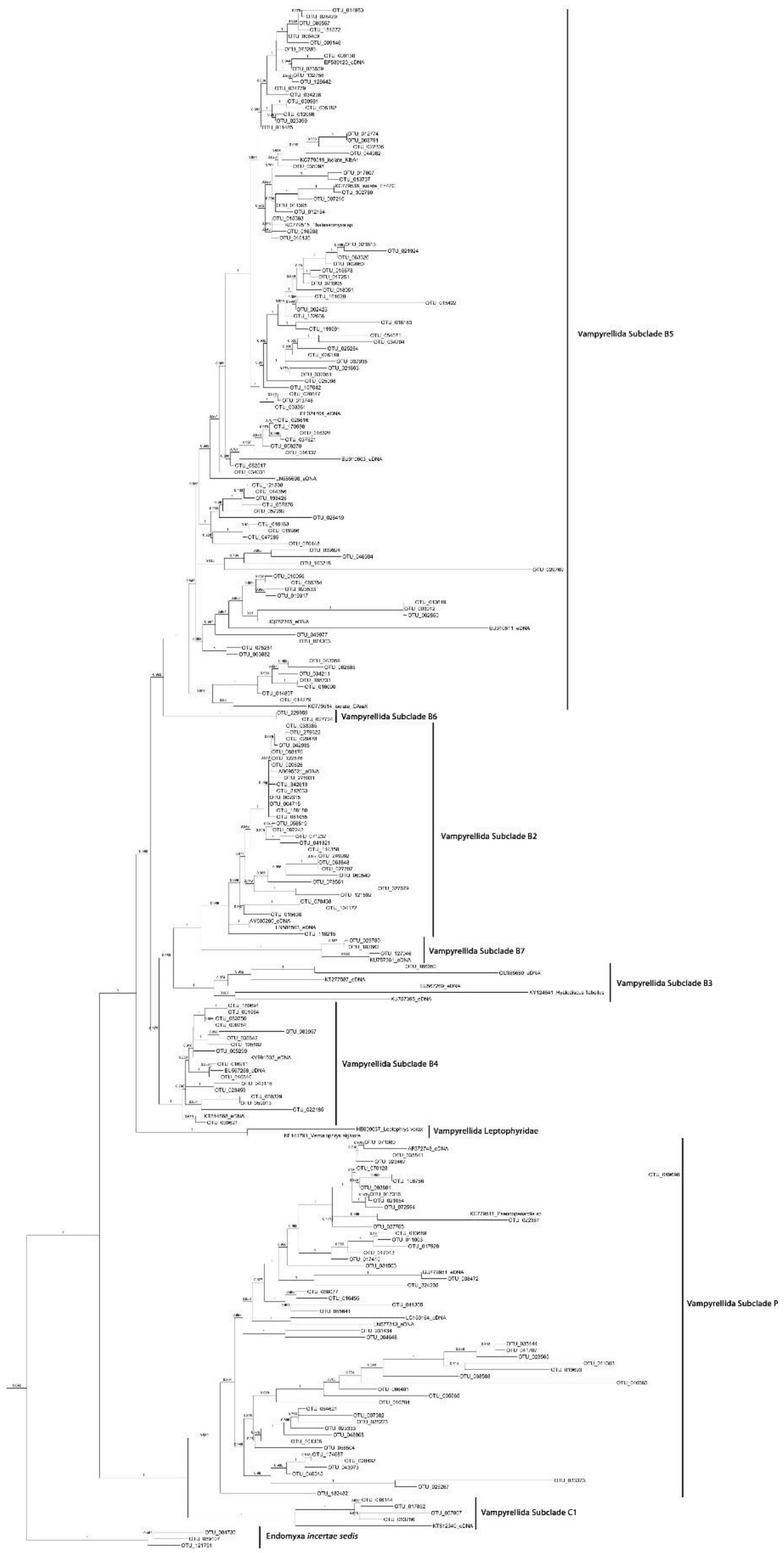


**Figure S2: PhyML tree of Vamyprellida** showing a detailed branching of the OTUs with previously published sequences (black dots) The OTUs recovered in this study showed a very similar pattern than was found in a previous study on the *BioMarKs* dataset using different libraries (Berney *et al.*, 2013).

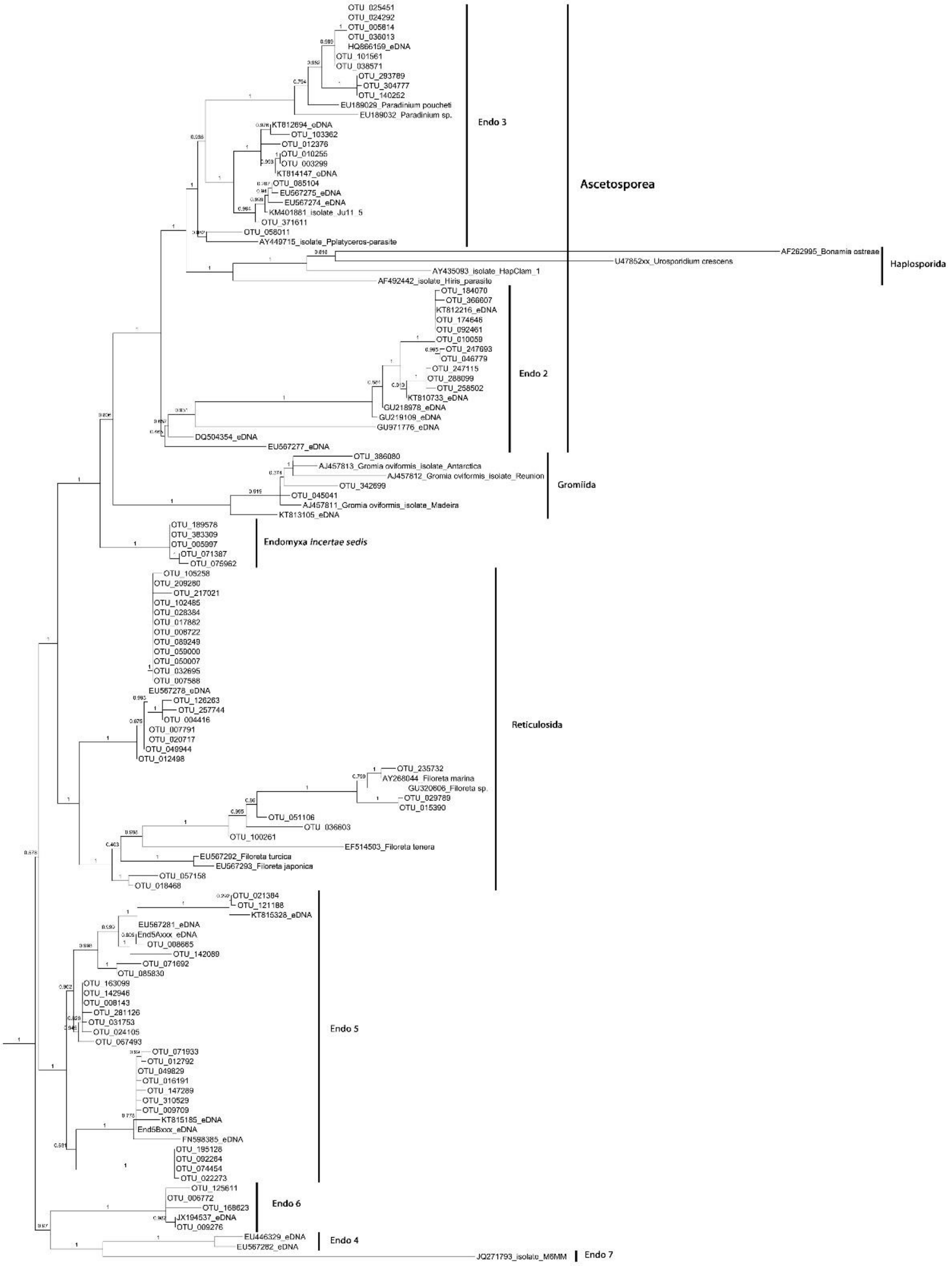


**Figure S3: PhyML tree Ascetosporea, Reticulosida and Gromiidia** together with the environmental clades Endo 4 to Endo 7 showing a detailed branching of the OTUs with previously published sequences (black dots). OTUs belonging to all major lineages but the Haplosporidia and the eDNA clades Endo4 and Endo7 were found.

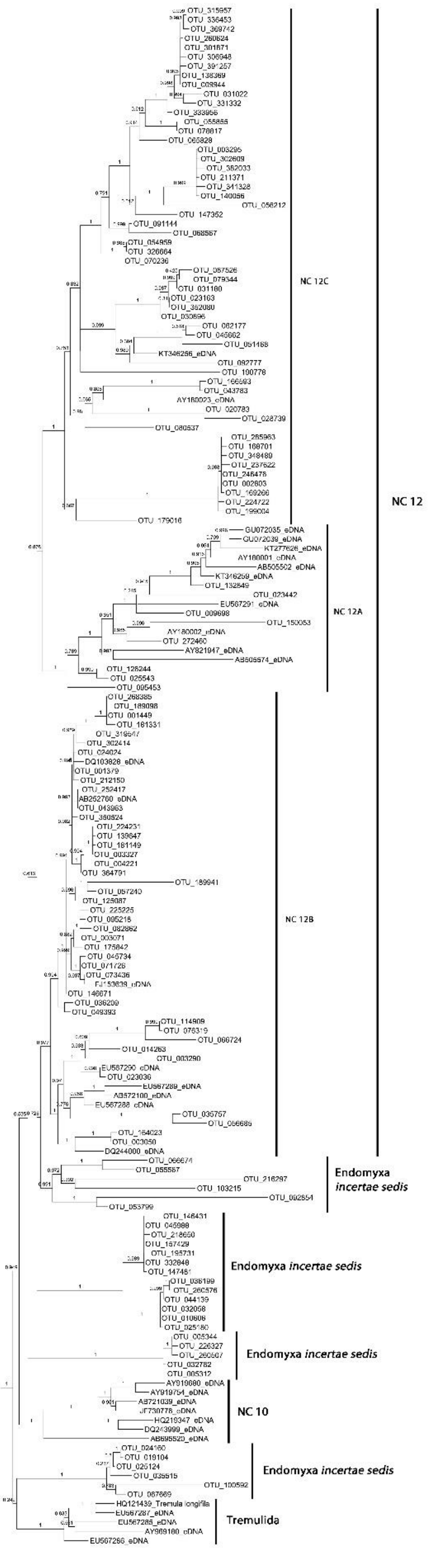


**Figure S4: PhyML tree of Tremulia, Aquavolon and the environmental clade NC12** showing a detailed branching of the OTUs with previously published sequences (black dots). Overall a huge diversity around the eDNA sequences that have been used to establish NC12 could be recovered. A more detailed study of this group will be needed to identify well supported new lineages.

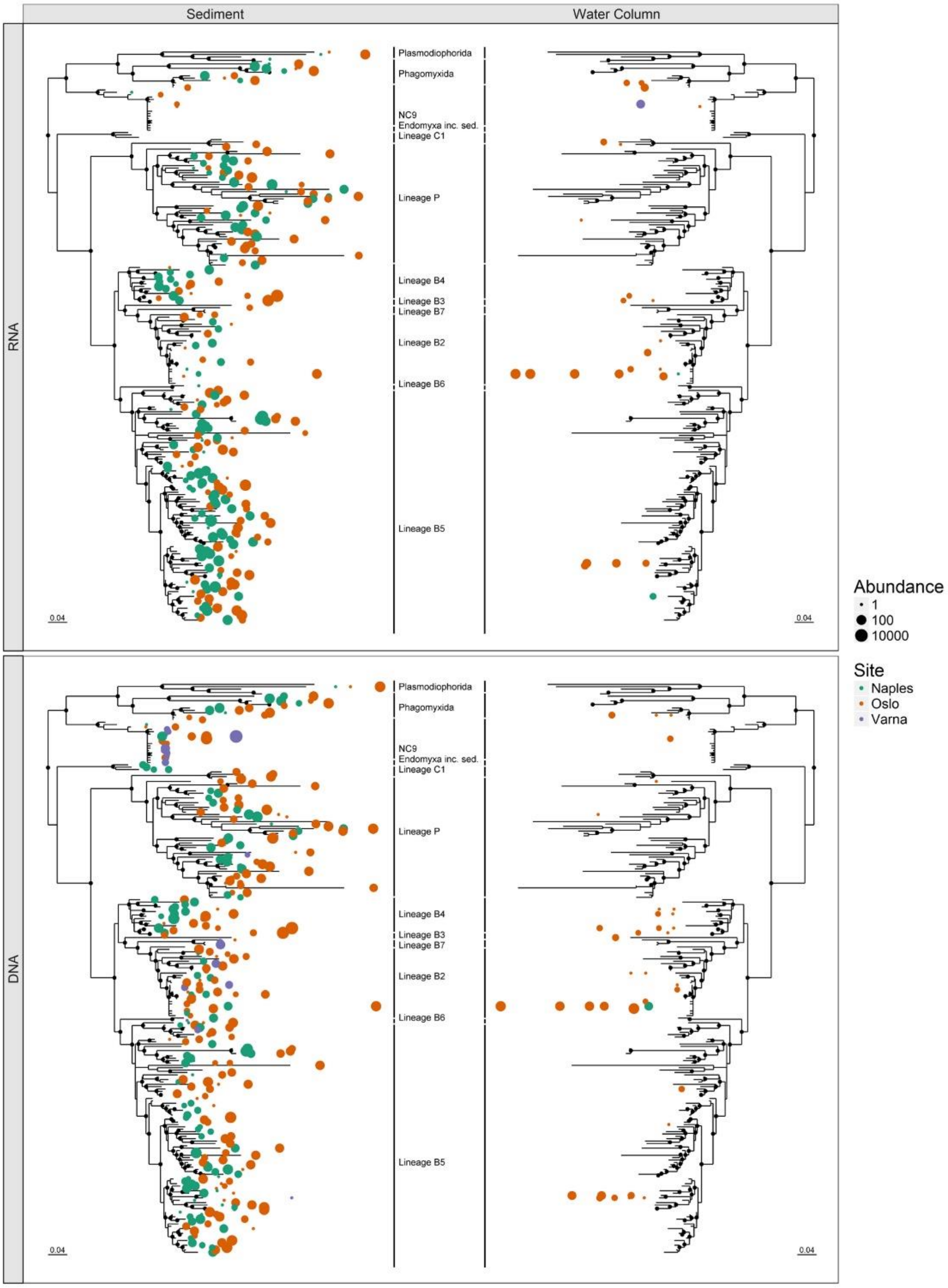


**Figure S5: Abundance (read numbers) and frequency (individual libraries where they were detected) of OTUs belonging to Phytomyxea and Vampyrellida.** Each sample that contained a respective OTU is shown as an individual dot, while the size indicates the number of reads that were recovered from each library. Black dots at the branching points of the tree indicate $\chi^2$ values above 0.95 from the PhyML analyses. Vampyrellida OTUs were evenly distributed in DNA and RNA samples from Varna and Oslo sediments, with some OTUs, mainly belonging to Vampyrellida Subclade B2 and B5 being abundant in the Water Column samples. Only few Vampyrellida OTUs were found in the samples from Varna. OTUs belonging to the Phytomyxea were mainly found in sediment samples with the notable exception of one NC9 OTU, which was found in high read numbers in the water column RNA sample from Varna. Trees were generated using the R-package 'phyloseq'.

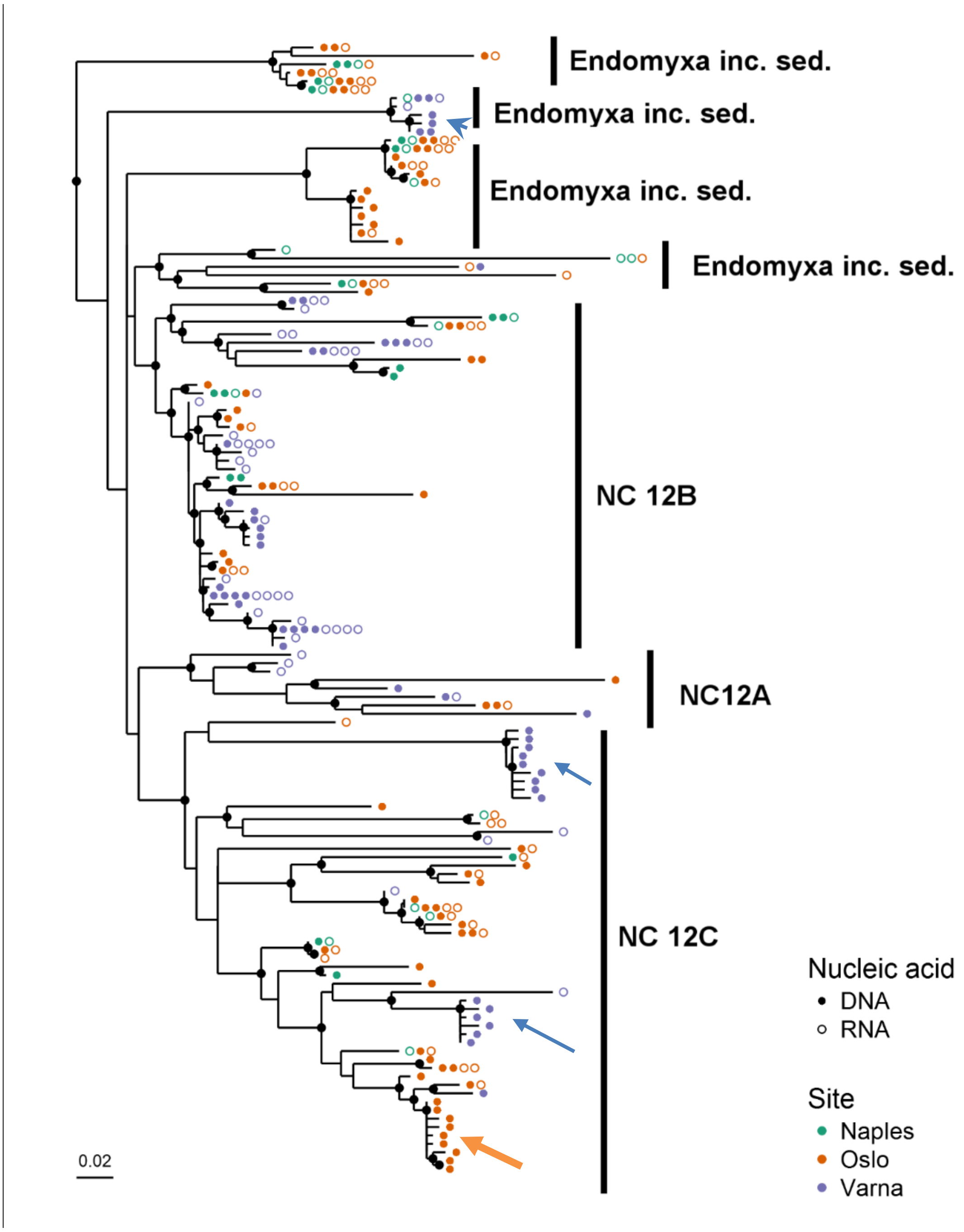


**Figure S6: Incidence of OTUs belonging to NC12.** Two long branching sub-clades of NC12 were found only in Varna DNA samples (blue arrows), as was one of the clades *incertae sedis* (blue arrowhead, basal to NC12). Another clade was found mainly in samples collected at Oslo (orange arrow). All the other sub-clades of NC12 showed a more even distribution of the OTUs across multiple samples and sampling sites. Each sample that contained a respective OTU is shown as an individual symbol, while dots denote DNA samples and circles represent RNA samples. Black dots at the branching points of the tree indicate $\chi^2$ values above 0.95 from the PhyML analyses. Tree was generated using the R-package ‘phyloseq’.

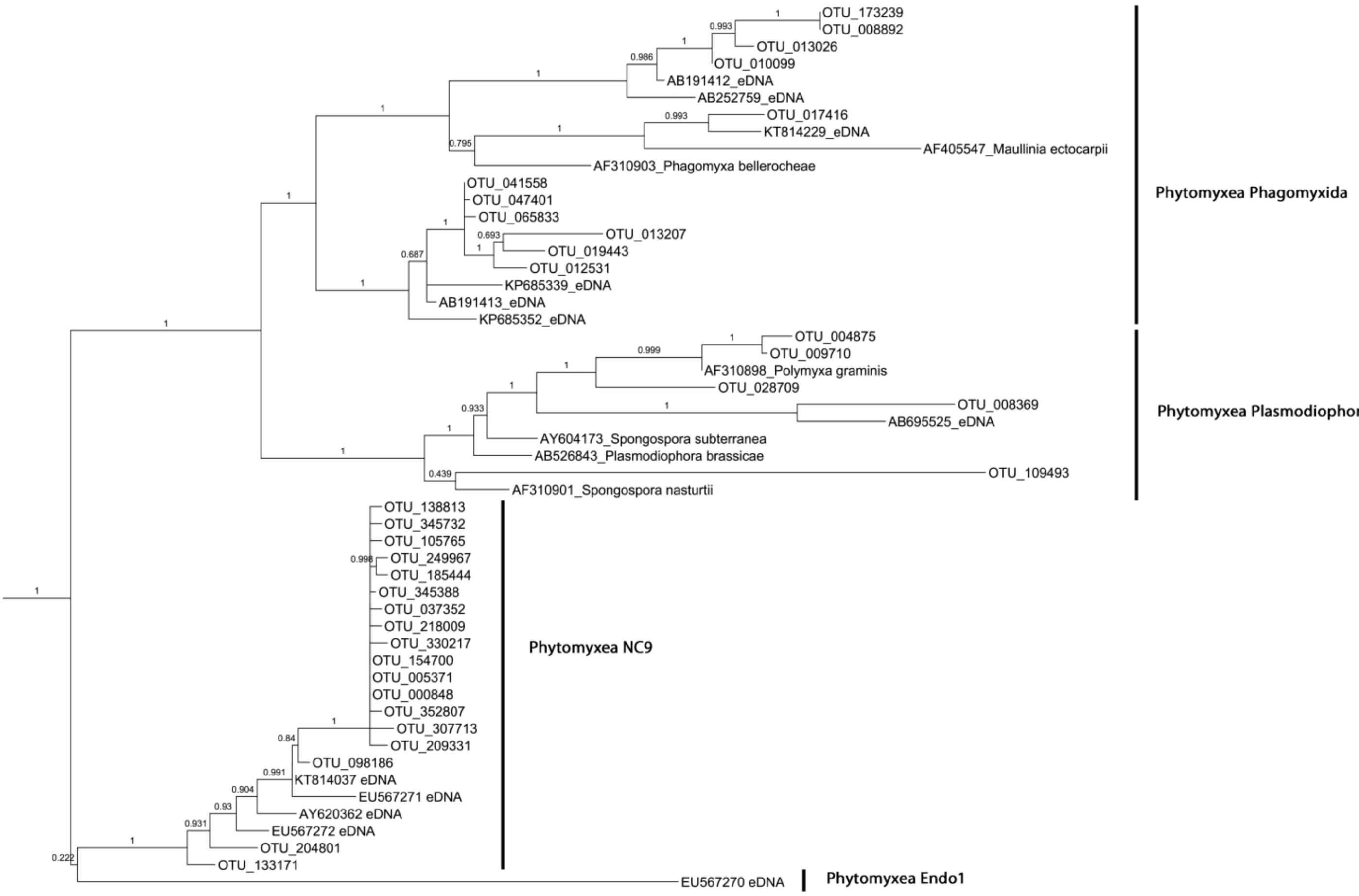


**Figure S7: PhyML tree of Phytomyxea** showing a detailed branching of the OTUs with previously published sequences (black dots). OTUs belonging to the marine Phagomyxida and the marine environmental Novel Clade 9 (NC9) were found. Also OTUs belonging to the soil and freshwater inhabiting Plasmodiophorida were detected. If these sequences belong to true marine Plasmdodiophorida needs to be determined in future studies.

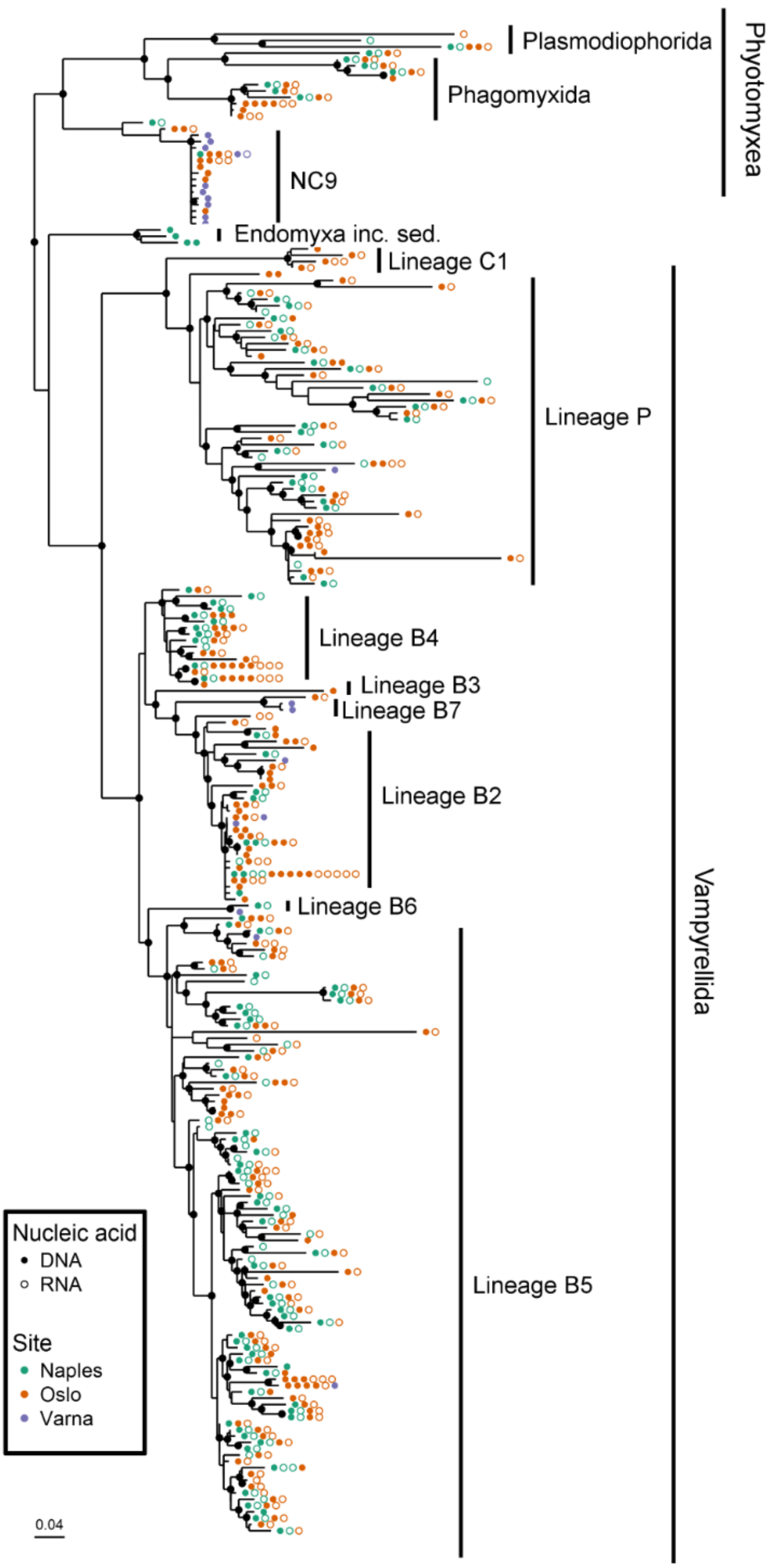


**Figure S8: Incidence2S6 of OTUs belonging to Phytomyxea and Vampyrellida.** Both groups showed a relatively even distribution of the OTUs across multiple samples and sampling sites. Each sample that contained a respective OTU is shown as an individual symbol, while dots denote DNA samples and circles represent RNA samples. Black dots at the branching points of the tree indicate $\chi^2$ values above 0.95 from the PhyML analyses. Tree was generated using the R-package 'phyloseq'.

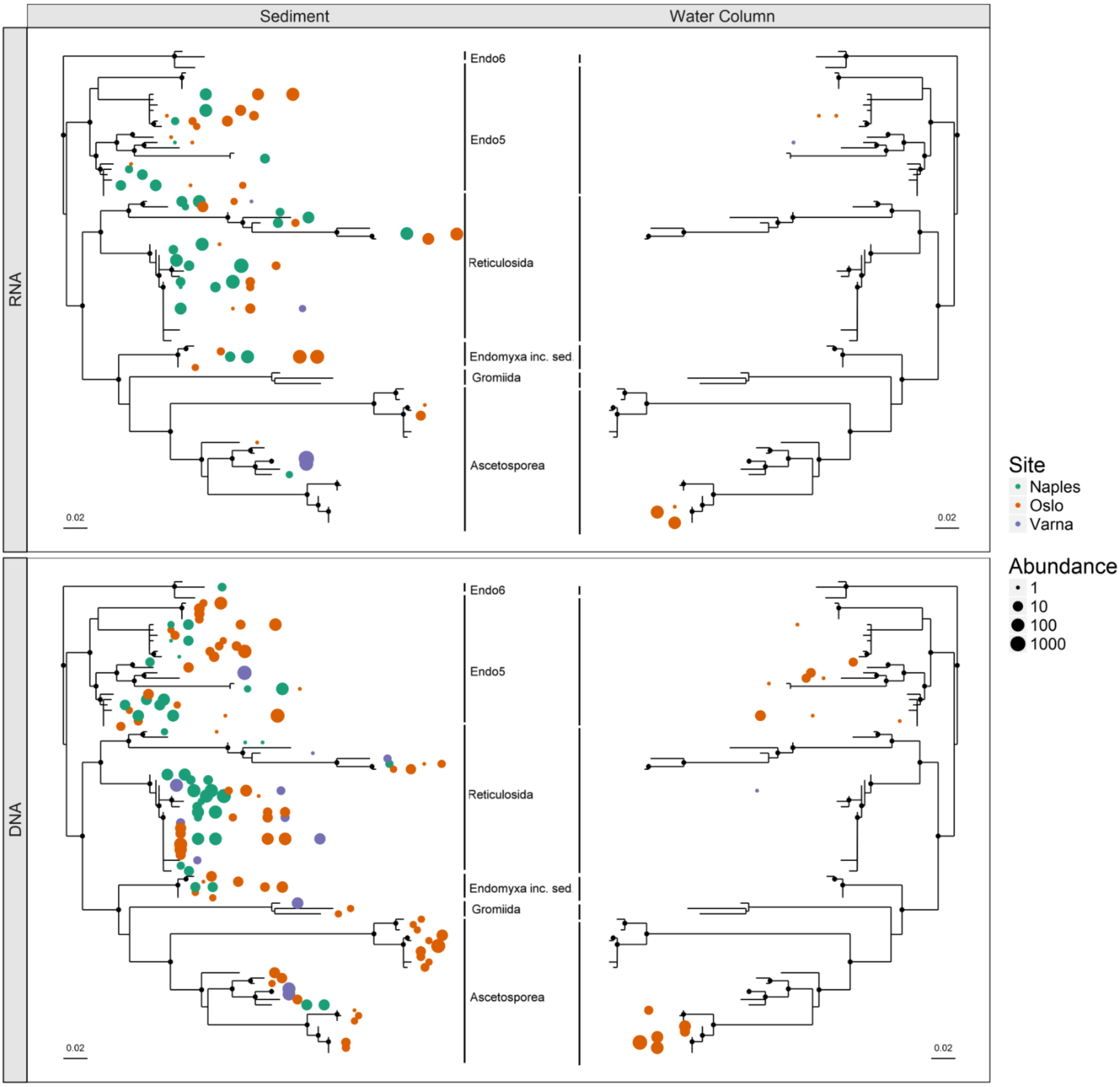


**Figure S9: Abundance (read numbers) and frequency (libraries where they were detected) of OTUs belonging to the Reticulosida, Gromiidia, the ascetosporean** clades Endo2 and Endo3 as well as the environmental clades Endo5 and Endo6. Each sample that contained a respective OTU is shown as an individual symbol, while the size indicates the number of reads that were recovered from each library. Black dots at the branching points of the tree indicate $\chi^2$ values above 0.95 from the PhyML analyses. The diversity of all groups is high in the sediment samples, with most clades being well represented at all three sampling sites. In the water column samples from Oslo a group of OTUs belonging to Endo3 could be detected in high read numbers. Trees were generated using the R-package ‘phyloseq’.

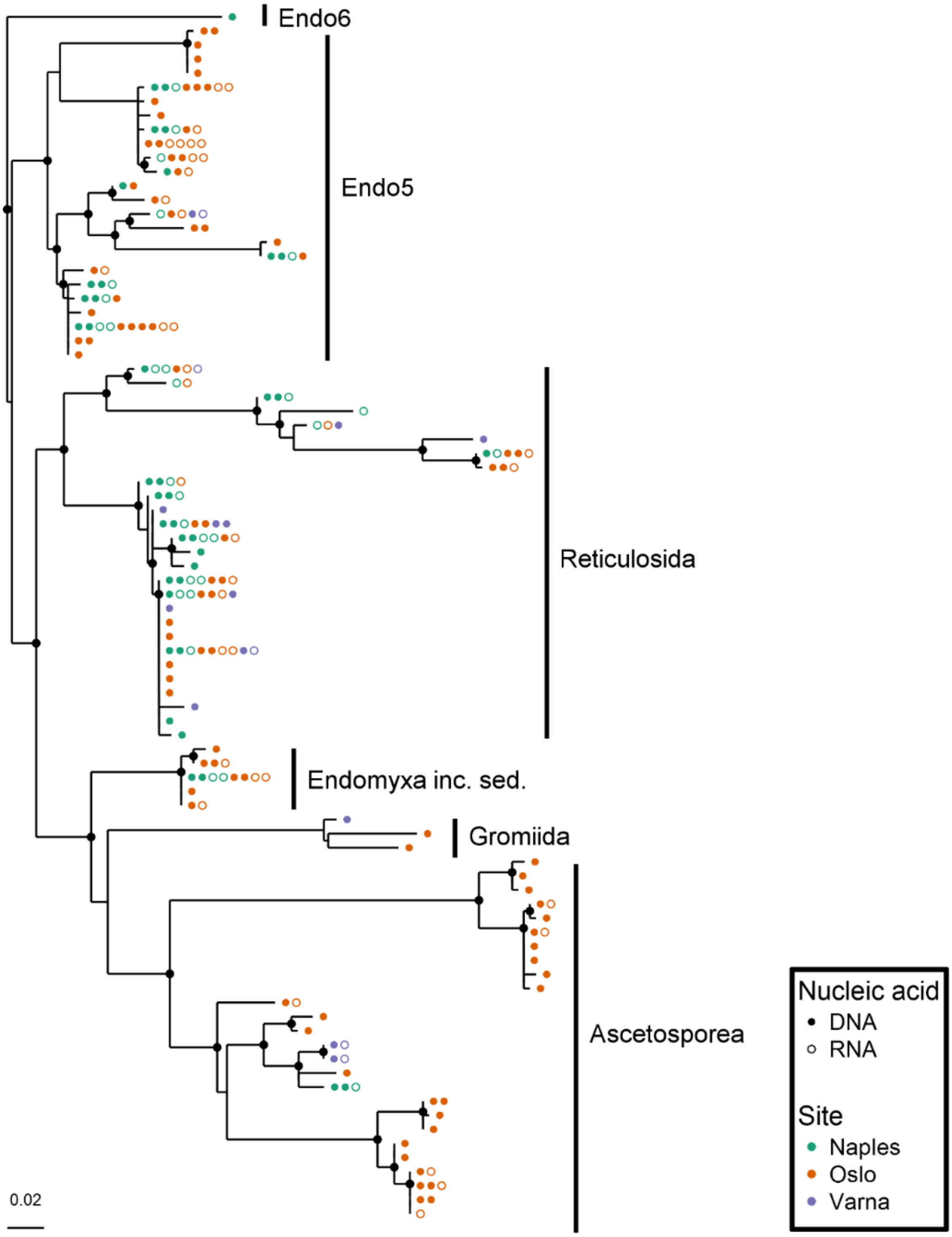


**Figure S10: Incidence of OTUs belonging to Reticulosida, Gromiida, Ascetosporea, and the environmental groups Endo5 and Endo6**. OTUs belonging to the Ascetosporea were nearly exclusively found in samples from Oslo while Reticulosida and Endo5 OTUs were more evenly distributed. Each sample that contained a respective OTU is shown as an individual symbol, while dots denote DNA samples and circles represent RNA samples. Black dots at the branching points of the tree indicate $\chi^2$ values above 0.95 from the PhyML analyses. Tree was generated using the R-package 'phyloseq'.
