## Supplementary material for "Unveiling hidden eukaryotes: diversity of Endomyxa (Rhizaria) in coastal marine habitats": Table S1 - Table S4

### *Unveiling hidden eukaryotes: the elusive Endomyxa (Rhizaria) are rich in function and diversity in coastal marine habitats*

#### *Supplementary Information - Tables*

Stefan Ciaghi[1,$], Cédric Berney[2,$], Sarah Romac[2], Frédéric Mahé[3], Colomban de Vargas[2], Olivier Jaillon[4], Martin Kirchmair[1], David Bass[5,6,*], Sigrid Neuhauser[1,*]

[1] Institute of Microbiology, University of Innsbruck, Technikerstr. 25, 6020 Innsbruck, Austria

[2] Sorbonne Université & CNRS, UMR 7144 (AD2M), Station Biologique de Roscoff, Place Georges Teissier, 29680 Roscoff, France

[3] CIRAD, UMR LSTM, Campus international de Baillarguet, 34398 Montpellier Cedex 5, France

[4] GENOSCOPE, Commissariat à l'Energie Atomique et aux Energies Alternatives (CEA), Institut de Génomique, 2 rue Gaston Crémieux, 91000 Evry, France

[5] Division of Genomics and Microbial Diversity, Dept of Life Sciences, Natural History Museum London, Cromwell Road, SW7 5BD, UK

[6] Centre for Environment, Fisheries and Aquaculture Science (Cefas), Barrack Road, The Nothe, Weymouth, DT4 8UB, UK

<u>**Supplementary Tables S1- S8**</u>

**Table S1:** Endomyxa OTUs generated using Endomyxa-specific and universal eukaryotic primers. **2**
**Table S2:** Read numbers and OTUs sorted by taxon and library. **3**
**Table S3:** Primers used to generate the 454 libraries. **4**
**Table S4:** Samples analysed **5**
**Table S5:** Multiplexing Tags (MIDs) **6**
**Table S6:** Genbank accession numbers **7**

**Table S1: Endomyxa OTUs** generated using Endomyxa-specific and universal eukaryotic primers. Endomyxa diversity was better represented using specific primers. Universal eukaryotic primers resulted in a bias towards Vampyrellida sequences. The three most abundant taxa are underlined in each column.

| Taxonomy | Endomyxa primers | | Eukaryote primers | | Total | |
|---|---|---|---|---|---|---|
| | OTUs [#] | OTUs [%] | OTUs [#] | OTUs [%] | OTUs [#] | OTUs [%] |
| Ascetosporea | 26 | 9.6 | 1 | 0.4 | 26 | 5.7 |
| Endo5 | 24 | 8.8 | 14 | 5.0 | 24 | 5.2 |
| Endo6 | 0 | 0.0 | 1 | 0.4 | 1 | 0.2 |
| Gromiida | 3 | 1.1 | 0 | 0.0 | 3 | 0.7 |
| NClade12 | <u>90</u> | <u>33.1</u> | <u>37</u> | <u>13.1</u> | <u>104</u> | <u>22.7</u> |
| Phytomyxea | 17 | 6.3 | <u>18</u> | <u>6.4</u> | 31 | 6.8 |
| Reticulosida | 26 | 9.6 | 14 | 5.0 | 27 | 5.9 |
| Vampyrellida | <u>51</u> | <u>18.8</u> | <u>179</u> | <u>63.5</u> | <u>204</u> | <u>44.5</u> |
| Endomyxa *incertae sedis* | <u>35</u> | <u>12.9</u> | <u>18</u> | <u>6.4</u> | <u>38</u> | <u>8.3</u> |
| **Total** | **272** | **100** | **282** | **100** | **458** | **100** |

**Table S2:** Read numbers and OTUs sorted by taxon and library. Values printed as [total read numbers (OTUs)]. As OTUs can be found in more than one sample, the total OTU-numbers can be higher than the total number of observed OTUs.

| | Sample | Ascetosporea | Gromiida | Reticulosida | Endo5 | Endo6 | Phytomyxea | Vampyrellida | NClade12 | Endomyxa incertae sedis | Total |
|---|---|---|---|---|---|---|---|---|---|---|---|
| **Naples** | **NB_SED_DNA** | 18 (1) | 0 (0) | 454 (11) | 78 (7) | 0 (0) | 84 (1) | 34 (3) | 52 (7) | 35 (5) | **755 (35)** |
| | **NB_SED_RNA** | 0 (0) | 0 (0) | 38 (4) | 12 (1) | 0 (0) | 0 (0) | 1 (1) | 0 (0) | 15 (2) | **66 (8)** |
| | **NB_WC_DNA** | 0 (0) | 0 (0) | 0 (0) | 0 (0) | 0 (0) | 0 (0) | 23 (1) | 0 (0) | 0 (0) | **23 (1)** |
| | **NB_WC_RNA** | 0 (0) | 0 (0) | 0 (0) | 0 (0) | 0 (0) | 0 (0) | 5 (2) | 0 (0) | 2 (1) | **7 (3)** |
| | **NB348_DNA** | 20 (1) | 0 (0) | 748 (10) | 198 (7) | 4 (1) | 478 (9) | 3 994 (91) | 9 (4) | 69 (8) | **5 520 (131)** |
| | **NB348_RNA** | 2 (1) | 0 (0) | 1 230 (13) | 192 (8) | 0 (0) | 296 (9) | 11 992 (110) | 93 (8) | 220 (10) | **14 025 (168)** |
| **Oslo** | **OF_SED_DNA** | 419 (19) | 4 (2) | 320 (11) | 668 (16) | 0 (0) | 1 087 (8) | 335 (34) | 608 (38) | 282 (23) | **3 723 (151)** |
| | **OF_SED_RNA** | 11 (3) | 0 (0) | 19 (2) | 80 (6) | 0 (0) | 1 (1) | 22 (7) | 53 (13) | 370 (10) | **556 (42)** |
| | **OF_WC_DNA** | 825 (6) | 0 (0) | 0 (0) | 35 (6) | 0 (0) | 3 (1) | 361 (5) | 0 (0) | 0 (0) | **1 224 (18)** |
| | **OF_WC_RNA** | 179 (3) | 0 (0) | 0 (0) | 1 (1) | 0 (0) | 1 (1) | 41 (2) | 0 (0) | 1 (1) | **223 (8)** |
| | **OF011_DNA** | 0 (0) | 0 (0) | 0 (0) | 2 (2) | 0 (0) | 1 (1) | 92 (7) | 0 (0) | 0 (0) | **95 (10)** |
| | **OF011_RNA** | 0 (0) | 0 (0) | 0 (0) | 0 (0) | 0 (0) | 3 (1) | 140 (5) | 0 (0) | 0 (0) | **143 (6)** |
| | **OF018_DNA** | 0 (0) | 0 (0) | 0 (0) | 0 (0) | 0 (0) | 4 (1) | 204 (11) | 0 (0) | 0 (0) | **208 (12)** |
| | **OF018_RNA** | 0 (0) | 0 (0) | 0 (0) | 0 (0) | 0 (0) | 0 (0) | 95 (6) | 0 (0) | 0 (0) | **95 (6)** |
| | **OF229_DNA** | 0 (0) | 0 (0) | 114 (8) | 86 (8) | 0 (0) | 808 (14) | 17 118 (116) | 27 (10) | 188 (10) | **18 341 (166)** |
| | **OF229_RNA** | 0 (0) | 0 (0) | 113 (9) | 134 (7) | 0 (0) | 375 (12) | 10 911 (114) | 122 (13) | 394 (14) | **12 049 (169)** |
| | **OF233_DNA** | 0 (0) | 0 (0) | 0 (0) | 1 (1) | 0 (0) | 1 (1) | 39 (8) | 0 (0) | 0 (0) | **41 (10)** |
| | **OF233_RNA** | 0 (0) | 0 (0) | 0 (0) | 1 (1) | 0 (0) | 14 (2) | 149 (6) | 0 (0) | 3 (1) | **167 (10)** |
| **Varna** | **VA_SED_DNA** | 180 (2) | 30 (1) | 144 (8) | 396 (1) | 0 (0) | 11 079 (10) | 104 (9) | 7 453 (30) | 931 (5) | **20 317 (66)** |
| | **VA_SED_RNA** | 1 964 (2) | 0 (0) | 3 (2) | 0 (0) | 0 (0) | 0 (0) | 0 (0) | 13 198 (15) | 880 (2) | **16 045 (21)** |
| | **VA_WC_DNA** | 0 (0) | 0 (0) | 1 (1) | 0 (0) | 0 (0) | 0 (0) | 0 (0) | 1 (1) | 7 (2) | **9 (4)** |
| | **VA_WC_RNA** | 0 (0) | 0 (0) | 0 (0) | 1 (1) | 0 (0) | 24 (1) | 0 (0) | 3 (2) | 0 (0) | **28 (4)** |
| | **VA119_DNA** | 0 (0) | 0 (0) | 0 (0) | 0 (0) | 0 (0) | 0 (0) | 0 (0) | 397 (5) | 0 (0) | **397 (5)** |
| | **VA119_RNA** | 0 (0) | 0 (0) | 0 (0) | 0 (0) | 0 (0) | 0 (0) | 0 (0) | 4 506 (12) | 0 (0) | **4 506 (12)** |
| | **VA122_DNA** | 0 (0) | 0 (0) | 0 (0) | 0 (0) | 0 (0) | 0 (0) | 0 (0) | 171 (6) | 0 (0) | **171 (6)** |
| | **VA122_RNA** | 0 (0) | 0 (0) | 0 (0) | 0 (0) | 0 (0) | 0 (0) | 0 (0) | 421 (10) | 0 (0) | **421 (10)** |
| | **VA133_DNA** | 0 (0) | 0 (0) | 0 (0) | 0 (0) | 0 (0) | 0 (0) | 0 (0) | 13 (2) | 0 (0) | **13 (2)** |
| | **VA133_RNA** | 0 (0) | 0 (0) | 0 (0) | 0 (0) | 0 (0) | 0 (0) | 0 (0) | 12 (2) | 0 (0) | **12 (2)** |
| | **Total** | **3 618 (26)** | **34 (3)** | **3 184 (27)** | **1 885 (24)** | **4 (1)** | **14 259 (31)** | **45 660 (204)** | **27 139 (104)** | **3 397 (38)** | **99 180 (458)** |

**Table S3: Primers used to generate the 454 libraries.** Primers used for the first PCR (PCR1) are highlighted in blue, primers used for the nested PCR (PCR2) are highlighted in red. First round PCR was identical for all samples, but two different sets of nested PCRs were performed using each forward primer with the PCR2 R-Primer mix. The ratio of each primer within an individual primer mix is given ("Ratio").

| Primer | Sequence, 5' -> 3' | Ratio | Specificity |
|---|---|---|---|
| **PCR1 F** | **Primer used individually** | | |
| s6f | GAGGRNAAGYCTGGTGCCAGCASC | | Eukaryota |
| **PCR1 R** | **Primer mix** | | |
| EndoR0 | CGACTTCTCCTTCTAAATGATAAG | 1 | Endomyxa-biased |
| EndoR1 | CGACTTCTCCTTCTAARYRDTAWG | 1 | Endomyxa-biased |
| EndoR2 | CGACTTCTCCTTCTAARYGHYWWG | 1 | Endomyxa-biased |
| EndoR3 | CGACTTYTCCTTCTARATRDYAWG | 1 | Endomyxa-biased |
| **PCR2 F** | **Primers used individually** | | |
| V4f-End | GTGCCAGCAGCCGCGGTAAYA | | Endomyxa-biased |
| V4f-Euk | CCAGCASCCGCGGTAAYWCC | | Eukaryota |
| **PCR2 R** | **Primer mix** | | |
| s1256R-C0 | CACCACCCATAGAATCAAGAAAGATCTTCA | 16 | Endomyxa-biased |
| s1256R-48 | CACYAHCCATAGAATCAAGAAAGRKCTKCA | 4 | Endomyxa-biased |
| s1256R-12 | CACCAMCCAWAGAATCAAGAAAGATCTGCA | 1 | NC12-biased |
| s1256R-Gr | CACCACCCATAWAATCAAGWAAGAKCTKCA | 1 | *Gromia*-biased |
| s1256R-Ha | CACYATKCATAGAATCAWGAAAGAACTTBA | 2 | Haplosporida-biased |
| s1256R-Ph | CACYACCCATAGAATCAAGAAAGAGCTKCA | 2 | Phagomyxida-biased |
| s1256R-Pl | CACCACCGAAGTGATCAAGAAAGAKCTKCA | 1 | Plasmodiophorida-biased |
| s1256R-Re | CACCAMCCATMRAATCAAGAAAGATCTTCA | 1 | *Reticulamoeba*-biased |
| s1256R-Fi | CACCACCCAYAGAATCAAGAAAGRTCTTCA | 2 | *Filoreta*-biased |
| s12S6R-Va | CACYAYCCATAGAATCAAGAAAGATCKTCA | 2 | Vampyrellida-biased |

**Table S4: Samples analysed**, listing the *BioMarKs* sample ID ("Sample"), collection site ("Site"), the type of nucleic acid that had been extracted from the original sample ("Target"), the type of sample that was collected ("Depth"), the category these samples were put in this study ("Pool"), the sequencing platform used ("Platform"), and whether general eukaryote primers ("Eukaryotes") or an Endomyxa-specific primer set was used ("Endomyxa"). Primers are listed in Table S2, tags are listed in Table S3.

| Sample | Site | Target | Depth | Pool | Platform | Primer |
|---|---|---|---|---|---|---|
| **NB348_DNA** | Naples | DNA | Sediment | Sediment | Illumina | Eukaryotes |
| **NB348_RNA** | Naples | RNA | Sediment | Sediment | Illumina | Eukaryotes |
| **OF011_DNA** | Oslo | DNA | DCM | Water Column | Illumina | Eukaryotes |
| **OF011_RNA** | Oslo | RNA | DCM | Water Column | Illumina | Eukaryotes |
| **OF018_DNA** | Oslo | DNA | DCM | Water Column | Illumina | Eukaryotes |
| **OF018_RNA** | Oslo | RNA | DCM | Water Column | Illumina | Eukaryotes |
| **OF229_DNA** | Oslo | DNA | Sediment | Sediment | Illumina | Eukaryotes |
| **OF229_RNA** | Oslo | RNA | Sediment | Sediment | Illumina | Eukaryotes |
| **OF233_DNA** | Oslo | DNA | DCM | Water Column | Illumina | Eukaryotes |
| **OF233_RNA** | Oslo | RNA | DCM | Water Column | Illumina | Eukaryotes |
| **VA119_DNA** | Varna | DNA | Anoxic | Sediment | Illumina | Eukaryotes |
| **VA119_RNA** | Varna | RNA | Anoxic | Sediment | Illumina | Eukaryotes |
| **VA122_DNA** | Varna | DNA | Anoxic | Sediment | Illumina | Eukaryotes |
| **VA122_RNA** | Varna | RNA | Anoxic | Sediment | Illumina | Eukaryotes |
| **VA133_DNA** | Varna | DNA | Anoxic | Sediment | Illumina | Eukaryotes |
| **VA133_RNA** | Varna | RNA | Anoxic | Sediment | Illumina | Eukaryotes |
| **NB_WC_DNA** | Naples | DNA | Water Column | Water Column | 454 | Endomyxa |
| **NB_WC_RNA** | Naples | RNA | Water Column | Water Column | 454 | Endomyxa |
| **NB_SED_RNA** | Naples | RNA | Sediment | Sediment | 454 | Endomyxa |
| **NB_SED_DNA** | Naples | DNA | Sediment | Sediment | 454 | Endomyxa |
| **OF_WC_DNA** | Oslo | DNA | Water Column | Water Column | 454 | Endomyxa |
| **OF_WC_RNA** | Oslo | RNA | Water Column | Water Column | 454 | Endomyxa |
| **OF_SED_RNA** | Oslo | RNA | Sediment | Sediment | 454 | Endomyxa |
| **OF_SED_DNA** | Oslo | DNA | Sediment | Sediment | 454 | Endomyxa |
| **VA_WC_DNA** | Varna | DNA | Water Column | Water Column | 454 | Endomyxa |
| **VA_WC_RNA** | Varna | RNA | Water Column | Water Column | 454 | Endomyxa |
| **VA_SED_RNA** | Varna | RNA | Sediment | Sediment | 454 | Endomyxa |
| **VA_SED_DNA** | Varna | DNA | Sediment | Sediment | 454 | Endomyxa |

**Table S5: Multiplexing Tags (MIDs)** and the samples they were used for. Each MID was used for the same sample type across the different sampling sites to even out bias within these sample pools. The column "sequencing" additionally states the sequencing depth on the 454 platform for the sample pool in the respective line.

| No MID | MID 1 | MID 2 | MID 3 | Sample | Sequencing |
|---|---|---|---|---|---|
| 0 | GTTGGTTG | CCTCCACA | AGTGGCAA | | |
| | | | | Naples | ¼ plate |
| DNA sediment | DNA water column | cDNA water column | cDNA sediment | Oslo | ¼ plate |
| | | | | Varna | ½ plate |

**Table S6: Genbank accession numbers** of sequences used for the phylogenetic placement alignment. Sequences from isolates and/or formally described sequences are printed in bold. In total 120 sequences were used, 29 of which were obtained from isolates according to their genbank metadata. Only 18 of these sequences are from formally described species.

| Genbank accession | Group | Sub-group | Species/Isolate |
|---|---|---|---|
| **HQ121439** | **Tremulida** | **Tremula** | ***Tremula longifila* strain ATCC50530** |
| **EU567285** | Tremulida | | clone db5 |
| **EU567286** | Tremulida | | clone lb6 |
| **EU567287** | Tremulida | | clone lb9 |
| **AY969180** | Tremulida | | clone dfmo4344061 |
| **AB695520** | *incertae sedis* | | clone MPE2-26 |
| **DQ243999** | Novel Clade 10 | | clone PCB07AU2004 |
| **HQ219347** | Novel Clade 10 | | clone AY2009C14 |
| **AY919680** | Novel Clade 10 | | clone LG01-12 |
| **AY919754** | Novel Clade 10 | | clone LG21-01 |
| **AB721039** | Novel Clade 10 | | clone RW15 2010 |
| **JF730776** | Novel Clade 10 | | clone Ch8A2mE7 |
| **AB505502** | Novel Clade 12 | NC12-A | clone RM1-SGM45 |
| **AB505574** | Novel Clade 12 | NC12-A | clone RM2-SGM66 |
| **AY180001** | Novel Clade 12 | NC12-A | clone CCA80 |
| **AY180002** | Novel Clade 12 | NC12-A | clone CCA33 |
| **AY821947** | Novel Clade 12 | NC12-A | clone CV1_B1_41 |
| **EU567291** | Novel Clade 12 | NC12-A | clone DB-2305-33 |
| **GU072035** | Novel Clade 12 | NC12-A | clone G512C1 |
| **GU072039** | Novel Clade 12 | NC12-A | clone G512H1 |
| **KT277626** | Novel Clade 12 | NC12-A | clone T12-D-12 |
| **KT346259** | Novel Clade 12 | NC12-A | clone HR_NC36_9 |
| **AB252760** | Novel Clade 12 | NC12-B | clone NAMAKO-20 |
| **AB572100** | Novel Clade 12 | NC12-B | clone ABC2_F09 |
| **DQ103828** | Novel Clade 12 | NC12-B | clone M2_18E01 |
| **DQ244000** | Novel Clade 12 | NC12-B | clone PCC4AU2004 |
| **EU567288** | Novel Clade 12 | NC12-B | clone DB-2305-21 |
| **EU567289** | Novel Clade 12 | NC12-B | clone DB-2703-18 |
| **EU567290** | Novel Clade 12 | NC12-B | clone DB-2703-19 |
| **FJ153639** | Novel Clade 12 | NC12-B | clone GoC1_F06 |
| **AY180023** | Novel Clade 12 | NC12-C | clone CCW46 |
| **KT346256** | Novel Clade 12 | NC12-C | clone HR_NC03_6 |
| **JQ271793** | *incertae sedis* | | **isolate M6MM** |
| **EU446329** | **lineage** Endo4 | | clone UI12B05 |
| **EU567282** | **lineage** Endo4 | | clone DB-2305-6 |
| **End5Axxx** | **lineage** Endo5 | | newly generated in this study |
| **EU567281** | **lineage** Endo5 | | clone sm5 |
| **KT815328** | **lineage** Endo5 | | clone 34c_10165 |
| **End5Bxxx** | **lineage** Endo5 | | newly generated in this study |
| **KT815185** | **lineage** Endo5 | | clone 21c_373 |

| Genbank accession | Group | Sub-group | Species/Isolate |
|---|---|---|---|
| **FN598385** | **lineage** Endo5 | | clone BS19_B6 |
| **JX194537** | *incertae sedis* | | clone 1301A09sw_184 |
| **AY268044** | **Reticulosida** | ***Filoreta*** | ***Filoreta marina*** |
| **GU320606** | **Reticulosida** | ***Filoreta*** | ***Filoreta sp.*** |
| **EU567293** | **Reticulosida** | ***Filoreta*** | ***Filoreta japonica*** |
| **EU567292** | **Reticulosida** | ***Filoreta*** | ***Filoreta turcica*** |
| **EF514503** | **Reticulosida** | ***Filoreta*** | ***Filoreta tenera*** |
| **EU567278** | Reticulosida | | clone DB-1412-C11 |
| **AJ457811** | **Gromiida** | ***Gromia*** | ***Gromia oviformis* isolate Madeira** |
| **AJ457812** | **Gromiida** | ***Gromia*** | ***Gromia oviformis* isolate Reunion** |
| **AJ457813** | **Gromiida** | ***Gromia*** | ***Gromia oviformis* isolate Antarctica** |
| **KT813105** | Gromiida | | clone 510c_58594 |
| **DQ504354** | Ascetosporea | **lineage** Endo2 | clone LC1043EP36 |
| **EU567277** | Ascetosporea | **lineage** Endo2 | clone DB-2305-52 |
| **GU971776** | Ascetosporea | **lineage** Endo2 | clone 1H2dD12 |
| **GU218978** | Ascetosporea | **lineage** Endo2 | clone GO_823_CERC |
| **GU219109** | Ascetosporea | **lineage** Endo2 | clone GW_1103_CERC |
| **KT810733** | Ascetosporea | **lineage** Endo2 | clone A7_70159 |
| **KT812216** | Ascetosporea | **lineage** Endo2 | clone 33c_98213 |
| **EU189029** | **Ascetosporea** | **Endo3 (Paradiniida)** | ***Paradinium poucheti*** |
| **EU189032** | **Ascetosporea** | **Endo3 (Paradiniida)** | ***Paradinium* sp. isolate PaEu41** |
| **HQ866159** | Ascetosporea | Endo3 **(Paradiniida)** | clone SGSP482 |
| **AY449715** | **Ascetosporea** | **Endo3 (Paradiniida)** | **ascetosporean isolate from *P. platyceros*** |
| **EU567274** | Ascetosporea | Endo3 **(Paradiniida)** | clone se16b |
| **EU567275** | Ascetosporea | Endo3 **(Paradiniida)** | clone DB-2305-51 |
| **KM401881** | Ascetosporea | Endo3 **(Paradiniida)** | **ascetosporean** isolate Ju11 5 |
| **KT812694** | Ascetosporea | Endo3 **(Paradiniida)** | clone A7 33426 |
| **KT814147** | Ascetosporea | Endo3 **(Paradiniida)** | clone 46c 60232 |
| **AF492442** | **Ascetosporea** | | **ascetosporean isolate from *H. iris*** |
| **AY435093** | Ascetosporea | Haplosporida | **ascetosporean** isolate from clam |
| **U47852xx** | **Ascetosporea** | **Haplosporida** | ***Urosporidium crescens*** |
| **AF262995** | **Ascetosporea** | **Haplosporida** | ***Bonamia ostreae*** |
| **EU567270** | Phytomyxea | **lineage** Endo1 | clone vm17 |
| **AY620362** | Phytomyxea | Novel Clade 9 | clone 7.4-5 |
| **EU567271** | Phytomyxea | Novel Clade 9 | clone jj9 |
| **EU567272** | Phytomyxea | Novel Clade 9 | clone nh6 |
| **KT814037** | Phytomyxea | Novel Clade 9 | clone 31c_102648 |
| **AB526843** | **Phytomyxea** | **Plasmodiophorida** | ***Plasmodiophora brassicae* isolate NGY** |
| **AF310898** | **Phytomyxea** | **Plasmodiophorida** | ***Polymyxa graminis*** |
| **AF310901** | **Phytomyxea** | **Plasmodiophorida** | ***Spongospora nasturtii*** |
| **AY604173** | **Phytomyxea** | **Plasmodiophorida** | ***Spongospora subterranea*** |
| **AB695525** | Phytomyxea | Plasmodiophorida | clone MPE2-31 |
| **AB191413** | Phytomyxea | *incertae sedis* | clone TAGIRI-5 |
| **KP685339** | Phytomyxea | Phagomyxida | clone F.13 2126 |
| **KP685352** | Phytomyxea | Phagomyxida | clone F.13 2073 |

| Genbank accession | Group | Sub-group | Species/Isolate |
|---|---|---|---|
| **AF310903** | **Phytomyxea** | **Phagomyxida** | ***Phagomyxa bellerocheae*** |
| **AF405547** | **Phytomyxea** | **Phagomyxida** | ***Maullinia ectocarpii* strain CCAP1538/1** |
| **KT814229** | Phytomyxea | Phagomyxida | clone 32c 2824 |
| **AB191412** | Phytomyxea | Phagomyxida | clone TAGIRI-4 |
| **AB252759** | Phytomyxea | Phagomyxida | clone NAMAKO-19 |
| **KT812340** | Vampyrellida | lineage C1 | clone 53c_50690 |
| **KC779511** | **Vampyrellida** | **subclade P** | **vampyrellid isolate NVam1** |
| **AF372743** | Vampyrellida | subclade P | clone LEMD004 |
| **GU479951** | Vampyrellida | subclade P | clone PR4 3E 90 |
| **LN577313** | Vampyrellida | subclade P | clone SICA1091 N11D3 18S E |
| **LC150164** | Vampyrellida | subclade P | clone B21 ek 24 |
| **HE609037** | **Vampyrellida** | **Leptophryidae** | ***Leptophrys vorax* strain LV.02** |
| **KF141791** | **Vampyrellida** | **Leptophryidae** | ***Vernalophrys algivore* isolate 20120808-63** |
| **KU757361** | Vampyrellida | lineage B7 | clone Plate2-35-M13R A04.ab1 |
| **AB695521** | Vampyrellida | lineage B2 | clone MPE2-27 |
| **LN586503** | Vampyrellida | lineage B2 | clone SICS1186 N9D2 16S A |
| **AY605200** | Vampyrellida | lineage B2 | clone Sey055 |
| **EU567268** | Vampyrellida | lineage B4 | clone op14 |
| **KY991032** | Vampyrellida | lineage B4 | clone C3CL106 |
| **KT814868** | Vampyrellida | lineage B4 | clone A7 94076 |
| **KY124641** | **Vampyrellida** | **lineage B3** | ***Hyalodiscus flabellus* isolate Hfla01** |
| **EU567269** | Vampyrellida | lineage B3 | clone T11 |
| **GU385680** | Vampyrellida | lineage B3 | clone ME Euk FW80 |
| **KT277587** | Vampyrellida | lineage B3 | clone T6-D-5 |
| **KU757395** | Vampyrellida | lineage B3 | clone Plate1-19-M13R A02.ab1 |
| **KC779515** | **Vampyrellida** | **lineage B5** | ***Thalassomyxa* sp. isolate MVa1x** |
| **KC779519** | **Vampyrellida** | **lineage B5** | **thalassomyxid isolate KibAr** |
| **KC779516** | **Vampyrellida** | **lineage B5** | **thalassomyxid isolate En42C** |
| **EF539120** | Vampyrellida | lineage B5 | thalassomyxid clone MB07.8 |
| **EF024704** | Vampyrellida | lineage B5 | thalassomyxid clone Elev 18S 1191 |
| **KC779514** | Vampyrellida | lineage B5 | vampyrellid isolate CAraX |
| **EU910603** | Vampyrellida | lineage B5 | clone D40 |
| **EU910611** | Vampyrellida | lineage B5 | clone D47 |
| **JQ782345** | Vampyrellida | lineage B5 | clone HD3bt0.37 |
| **LN585698** | Vampyrellida | lineage B5 | clone SIHB.1.1044 N9D0 16S B |
